# Proximal tubule-enhanced kidney organoids with S3-like cell enrichment model physiological responses

**DOI:** 10.64898/2026.01.29.702488

**Authors:** Sophia Mah, Ker Sin Tan, Sean B. Wilson, Kynan T Lawlor, Beatrice Gabbin, Emma I. Scully, Marella Cuevas, Richard J Mills, Melissa H Little, Jessica M Vanslambrouck

**Author notes:** Author for correspondence: J.M.V. Equal first author contribution (Sophia Mah, Ker Sin Tan). Equal last author contribution (Melissa H Little, Jessica M Vanslambrouck).

## Abstract

Chronic kidney disease (CKD) is a major global health challenge affecting >840 million people, with rising prevalence driven by common conditions including diabetes, hypertension, obesity, and drug toxicity. The proximal tubule (PT) is central to kidney function yet is highly injury-prone, particularly within the S3 segment which plays key roles in CKD progression. Despite decades of *in vitro* PT model development, achieving accurate functional and structural segmentation of the PT with distinct PT sub-cell types has remained challenging.

Here, we report an enrichment of S3-like cells within PT-enhanced kidney organoids (PT-EKO) that confers segment-specific functionality. This advanced platform facilitated modelling of physiological responses to environmental changes, including glucose levels, fatty acid substrate availability, and fluid sheer stress. Integrating with both static and organ-on-chip culture systems, the translational potential of S3-enriched PT-EKOs was underscored by their amenability to scale-up via cryopreservation of day 13 progenitors with retained differentiation capacity. PT-EKO applications were further broadened as an expandable high-yield source of S3-like PT cells, retaining transporter functionality across multiple passages.

Together, these findings represent an advance towards more physiologically relevant, human-specific modelling of PT responses supporting therapeutic approaches for CKD, including drug development, toxicity evaluation and cellular therapy.

**SIGNIFICANCE STATEMENT:** Advancing stem cell-based proximal tubule models, here we identify proximal tubule-enhanced kidney organoids (PT-EKO) as an enriched source of the most injury-prone cell type in the nephron; S3 proximal tubule cells. This advanced platform provides physiologically-relevant modelling of tubular responses to changes in the metabolic and biophysical environment, as well as a scalable source of high-quality PT cells, paving the way for more accurate modelling, screening, and cellular therapy applications.

## INTRODUCTION

Chronic kidney disease (CKD) continues to represent a major underdiagnosed global health challenge attributed to 1.2 million deaths annually ^1^. Ranking among the leading causes of mortality and morbidity, approximately 700 million people are living with CKD worldwide, with this prevalence rising to >850 million with the inclusion of acute kidney injury (AKI) and kidney failure ^1^. The increasing incidence of CKD is driven by aging populations, increased incidence of diabetes, obesity, hypertension, and drug toxicity. Among these, diabetes mellitus accounts for 30– 50% of CKD cases ^2^. Prolonged hyperglycemia and excessive glucose filtration promote kidney damage through mechanisms such as glucose and albumin overload, metabolic stress, and lipotoxicity. However, the renal consequences of disease are often compounded by therapeutic interventions. Nephrotoxicity is a common consequence of medications, particularly in hospitalized patients, with 14% - 26% of AKI predicted to be drug-induced, predisposing patients to CKD progression ^3,4^.

At a cellular level, these conditions compromise the vulnerable functional units of the kidney known as nephrons. With around 1 million nephrons per kidney, these structures carry out the vital processes of blood filtration, nutrient reabsorption, waste excretion, and maintenance of body homeostasis. Within each nephron, it is the specialized early proximal tubule segment that is central to kidney function, reabsorbing the bulk of nutrients, electrolytes, and water, while simultaneously secreting wastes, generating energy and vitamins, and regulating blood pressure ^5^. However, its high metabolic activity and exposure to toxic stressors also makes the PT prone to injury, with this susceptibility changing along its length. While the early convoluted PT segment (comprised of S1–S2 cell types) performs the bulk of solute transport, it is the downstream straight PT segment / pas recta (comprised of S3 cell types) that is considered the Achilles heel of the PT. Possessing fewer mitochondria than the S1–S2, the hypoxic outer medulla that the S3 segment resides in and its predominant role in drug clearance makes S3 PT cells disproportionately susceptible to ischemic and toxic injury, as well as diseases such as diabetes ^6,7^.

Following injury, the S3 segment and PT overall has some capacity to repair through dedifferentiation and proliferation of surviving epithelial cells ^8,9^. However, the limited nature of this regenerative capacity results in progressive functional decline and CKD onset. As a result, approximately 4 million people worldwide rely on regular dialysis; a costly and burdensome treatment with significant quality of life impacts ^10^. For the >1 million patients living with end-stage renal disease, transplantation remains the best therapeutic option. However, donor organ supply falls drastically short of demand.

These challenges have intensified efforts to develop efficient, cost-effective, and human-relevant *in vitro* models of the PT. The last few decades have seen the exploration of a range of approaches, such as 2D static cultures of primary or immortalized PT cells ^11–16^, PT-like cells directly derived from stem cells or via reprogramming ^17–20^, self-organizing kidney organoids containing PT structures maintained under static or flow conditions ^21–30^, and PT cells cultured under flow as 3D tubes or within microfluidic channels ^31–38^. Despite these advances, current models are limited by limitations in transporter expression, expandability, and recapitulation of structural and functional PT segmentation ^39,40^. To date, recreation of the distinct cellular subtypes that define individual PT segments has been limited. This is particularly the case for the S3 segment, the most relevant to AKI / CKD-associated injury and an early target in conditions such as diabetes ^7^.

We previously reported the development of proximal tubule–enhanced kidney organoids (PT-EKOs) through optimized specification of nephron progenitors from human pluripotent stem cells (hPSCs) ^41^. PT-EKOs were the first model to display elongated proximal nephrons with more mature marker expression compared with standard organoid protocols, providing a superior platform for infectious disease modeling and drug toxicity studies ^41–43^.

Within this study, enrichment for S3-like cells is identified within PT-EKO, supported by transcriptomic signatures akin to adult human PT cells and protein evidence of PT subdivision. The presence of S3 segmentation enabled appropriate responses to metabolic substrate shifts. This was evidenced through activation of Kidney Injury Molecule-1 (KIM-1) ectodomain shedding in response to elevated glucose, as well as improved mitochondrial activity in response to long chain fatty acids and fluid sheer stress within organ-on-a-chip systems. The translation utility of this model was demonstrated by showing scalability via cryopreservation of PT-EKO progenitors and the capacity to isolate S3 PT cells with stable identity across multiple passages. Together, these findings define a robust, human-relevant platform for accurate modeling of S3 PT responses. This may pave the way to scalable drug discovery, cell-specific toxicity testing and cellular therapies to combat CKD.

## RESULTS

### PT-EKO possess distinct PT sub-segments and are enriched for S3-like cell types

We have previously demonstrated improved metanephric nephron progenitor identity through prolonged differentiation, deriving organoids (PT-EKO) with a high proportion of elongated PTs and broad PT marker expression (Figure 1A, Supplementary Figure 1A) ^41^. Given that structural PT subdivision underpins PT functionality (Figure 1B), comprehensive re-analyses were performed on PT-EKO single cell RNA sequencing (scRNAseq) datasets to evaluate the relative presence of distinct S1-S2 (convoluted PT) and S3 (straight PT) cell populations.

**Figure 1:**
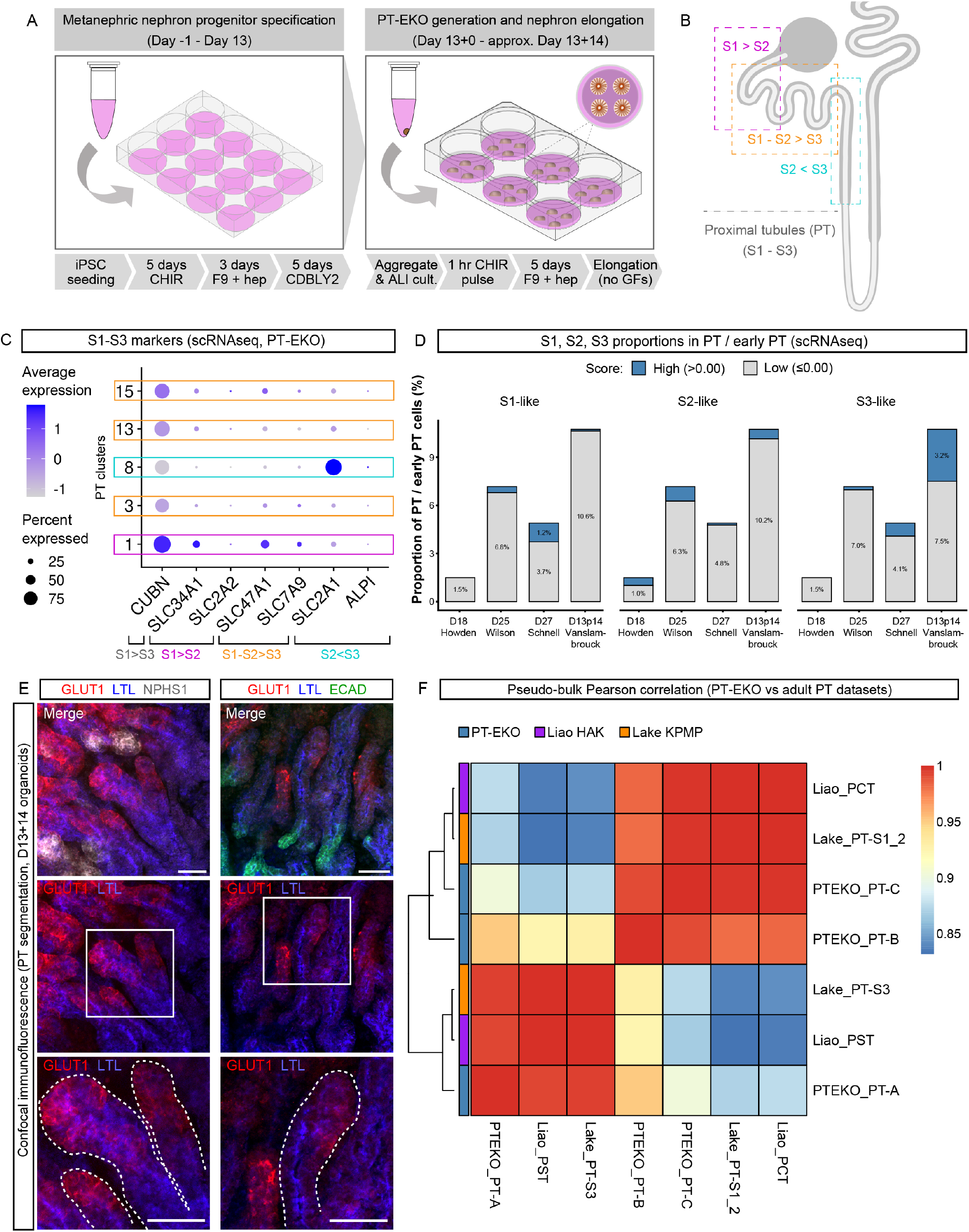
PT-EKOs possess distinct PT sub-segments and are enriched for S3-like cell types. **A.** Schematic depicting PT-EKO generation from monolayer differentiation (day 0 – day 13) to organoid generation and development (day 13+0 onwards). CHIR = CHIR99021; F9 = FGF9; hep = heparin; CDBLY2 = nephron progenitor maintenance media 2 ^43^; GFs = growth factors. **B.** Schematic of the nephron depicting proximal tubule (PT) segments and PT marker expression gradients (S1>S2 = higher in S1 segment; S1-S2>S3 = higher in S1-S2 segments, expression declining to S3; S2>S3 = higher in S2 segment). **C.** Dot plot depicting the expression of known PT segment-specific markers (S1-S3 [convoluted] and S3 [straight]) in PT clusters within the PT-EKO scRNAseq dataset, including 4 hashed biological replicates, merged. Dot colour and size in dot plots represent unscaled gene expression and percentage of cells expressing each gene, respectively. **D.** Bar chart depicting the percentage of cells scored as S1, S2, and S3 identities within the PT and early PT populations of published scRNAseq kidney organoid datasets (‘D18 Howden’ ^50^, ‘D25 Wilson’ ^51^),‘D27 Schnell’ ^27^, and ‘D13p14 Vanslambrouck’. Cells were deemed positive with a score above 0. **E.** Confocal immunofluorescence images of PT-EKOs stained with markers of proximal tubule brush border membrane, S1-S3 (*Lotus tetragonolobus* lectin [LTL]; blue), proximal tubule S2<S3 basolateral glucose transporter (GLUT1; red), podocytes of the glomeruli (NEPHRIN [NPHS1]; grey), and distal nephron epithelium (E-CADHERIN [ECAD]; green). Dotted lines highlight examples of LTL-positive lengths of PTs. Scale bars represent 50µm. **F.** Pseudo-bulk Person’s correlation heatmap, calculated on the pseudo-bulk transcriptomic profiles of PT-clusters from PT-EKO (PTEKO_PT-A [merged S3-like clusters 8 and 3]), PTEKO_PT-B [S1-S2-like cluster 15], and PTEKO_PT-C [merged S1-S2-like clusters 1 and 13]), compared to 2 adult human kidney reference datasets (Liao *et al.* [2020] and Lake *et al.* [2023, Kidney Cell Atlas] ^52,53^) containing S3 (Liao_PST and Lake_PT-S3) and S1/S2 identities (Liao_PCT and Lake_PT-S1_2).

Expression of *CUBN*, an S1>S3 marker (expressed S1 – S3, highest in S1 ^44^) suggested the presence of 5 PT clusters (clusters 1, 3, 8, 13, and 15), with highest expression observed in cluster 1, followed by 15 and 13 (Figure 1C). To more deeply investigate cluster identity, analyses were performed for a range of transporters known to show spatially distinct expression patterns along the PT of adult human kidney. Consistent with the known gradient of CUBN expression from S1>S3, this demonstrated highest expression of earlier S1>S2 PT segment markers (*SLC34A1, SLC2A2*) in clusters 1, 15, and 13 (Figure 1C). In contrast, marker *SLC2A1* (encoding the basolateral GLUT1 transporter more active in S3 cells ^45^), showed higher expression levels in a larger proportion of cells compared to S1-S2 markers overall and was largely restricted to cluster 8. A smaller proportion of these cells also expressed high levels of the more mature S3 marker, *ALPI*. In combination with low S1-S2 marker expression, these findings suggested this cluster represented cells more similar to the S3 portion of the PT (Figure 1C).

These results were strengthened by glucose transporter expression signatures. Glucose transport is one of the key roles of the PT, reabsorbing nearly 100% of filtered glucose via distinct pairs of transporters active in the early S1-S2 (apical SGLT2 [*SLC5A2*] / basolateral GLUT2 [*SLC2A2*]) and later S3 (apical SGLT1 [*SLC5A1*] / basolateral GLUT1 [*SLC2A1*]) segments ^46–49^. While expression of SGLT-encoding genes (*SLC5A1* and *SLC5A2*) were low overall, analyses of glucose transporters in merged earlier PT clusters (1, 3, 13, and 15) and the putative latest PT cluster (8) confirmed highest expression of *SLC2A1*/ *SLC5A1* in the later S3-like population and *SLC2A2* / *SLC5A2* in the earlier S1-S2>S3 populations as anticipated (Figure 1C, Supplementary Figure 1B), suggesting an enrichment for S3-like PT cells.

To further investigate potential S3 enrichment within PT-EKOs, the proportions of PT cell populations in day 13+14 (D13+14) PT-EKO were compared to published scRNAseq datasets using broader PT marker panels for S1-like (S*LC6A19, SLC36A2, RBP4, SLC2A2, SLC5A2, SLC34A1*), S2-like (*CA4, LYZ, SLC6A20, SLC15A2*) and S3-like (*SLC5A1, ALPI, SLC2A1, ALC6A20, SLC15A2*) (Figure 1D,

Supplementary Table 1). Comparisons were made to two standard Takasato protocol-derived kidney organoid datasets (D7+12 ‘D18 Howden’ ^50^ and D7+18 ‘D25 Wilson’ ^51^), and a recently published proximal-biased kidney organoid dataset, ‘D27 Schnell’ (Day27Ly29) ^27^. This demonstrated a higher proportion of PT and early PT cells with high S3-like PT scores within PT-EKO compared to standard and proximal-biased datasets (Figure 1D). S3 high-scoring cells were also present at higher proportions in PT-EKO compared to S1 and S2 high scoring populations, further suggesting an enrichment of this population within PT-EKO.

This was supported by immunofluorescence for the GLUT1 (*SLC2A1*) transporter, a high-affinity basolateral glucose transporter most active in later S2-S3 straight segments ^46^ (Figure 1E, Supplementary Figure 1C). Immunofluorescence for GLUT1 in combination with Lotus tetragonolobus lectin (LTL; marking brush border membrane along the length of the PT, S1-S3) revealed restriction of GLUT1 expression to only a portion of LTL-positive PTs (Figure 1E). Together with gene expression analyses, these results suggested structural separation of the PTs of PT-EKO, composed of S1-S2-like and S3-like cell types, with enrichment for S3 cells within the dataset.

To compare defined PT cell identities within PT-EKO to that of the adult PT, where cells have matured into distinct S1/S2/S3 subtypes, a more comprehensive PT segment-specific anchor gene panel curated from literature (prioritizing human data where available; Supplementary Table 1, Supplementary Figure 1) was used to further validate proposed PT identities and merge similar datasets into S1-S2-like and S3-like. Using this approach, clusters determined to be S3-like (clusters 3 and 8 [A]) and S1-S2-like (clusters 13 and 1 [C], and cluster 15 [B]) were merged to create ‘PTEKO_PT-A’ and ‘PTEKO_PT-B’ / ‘PTEKO_PT-C’ (Figure 1F; Supplementary Figure 1D). Datasets were pseudo-bulked to facilitate comparisons between PT populations of PT-EKO and publicly available human adult kidney reference datasets which had defined PT sub-populations (Liao *et al.* [2020] and Lake *et al.* [2023, Kidney Cell Atlas]). ^52,53^

Both adult reference datasets were confirmed to contain identifiable convoluted (S1-S2: ‘Liao_PCT’ and ‘Lake_PT-S1_2’) and straight (S3: ‘Liao_PST’ and ‘Lake_PT-S3’) PT populations (Figure 1F, Supplementary Figure 1D). Variability was noted in the expression level of individual PT segment anchor genes when comparing S1/S2 and S3 populations across all datasets, including between the adult reference datasets themselves (Liao and Lake) (Supplementary Figure 1D), highlighting the challenge of overlapping gene expression gradients through the PT length when defining such populations. However, despite these variations and large differences in maturity between adult and organoid models, the PTEKO_PT-A population (with lower S1/S2 markers and higher S3 markers) was found to correlate most highly with the S3-like adult datasets (Liao_PST and Lake_PT-S3) (Figure 1F). In agreement, Liao_PCT and Lake_PT-S1_2 showed highest correlation with PTEKO_PT-C (S1>S2-like), strengthening the identity of distinct PT cell subtypes in PT-EKO.

### PT-EKO show selective glucose transport activities and responses

We have previously shown that PT-EKO possess PT-specific transporter-dependent uptake of organic cations (4′,6-diamidino-2-phenylindole) and expected toxicity response following exposure to the nephrotoxic chemotherapeutic, cisplatin ^41^. To explore PT functionality beyond exogenous nephrotoxic stimuli, the response to physiological metabolic substances was assessed, including the activity of segment-specific PT glucose transporters such as SGLT2 (expressed on the apical membrane of earlier S1-S2 convoluted PT segments) and GLUT1 (high affinity basolateral glucose transporter most active in later S2-S3 straight segments) ^46^. Glucose transport in PT-EKO was assessed using the green-fluorescent D-glucose analogue, 2-[N-(7-nitrobenz-2-oxa-1,3-diazol-4-yl)amino]-2-deoxy-D-glucose (2-NBDG ^54^ ). Following 2 hours incubation in 2-NBDG, live confocal imaging revealed high levels of 2-NBBG uptake into PT-EKO (Figure 2A) compared to control PT-EKOs without substrate exposure (Supplementary Figure 2A).

**Figure 2:**
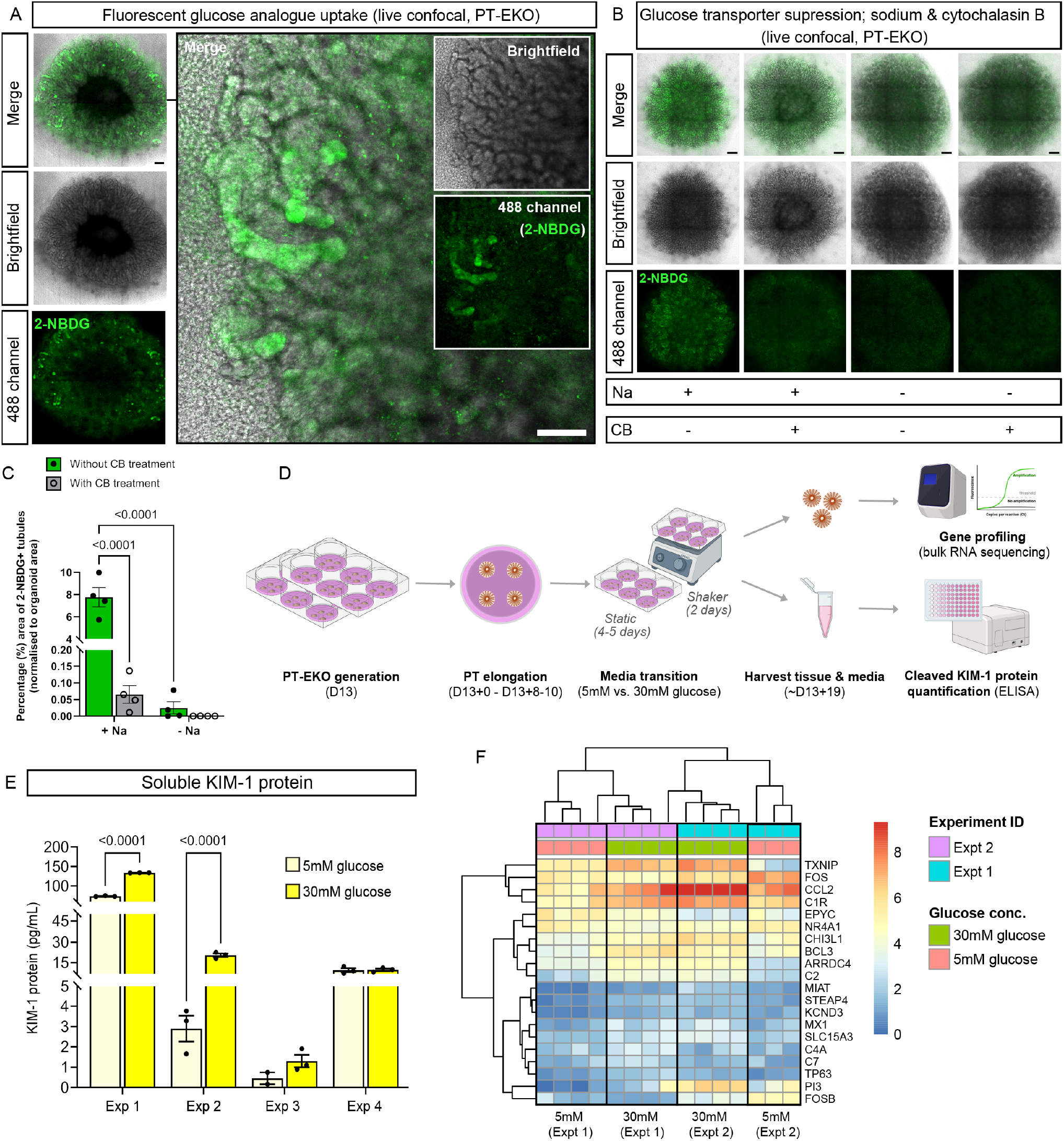
PT-EKOs display segment-specific PT transport and glucose responsiveness. **A-C**. Live confocal imaging and quantification of D13+14 PT-EKO incubated in either green fluorescent glucose analogue (2-NBDG) alone ([**A**]; right panel depicts higher magnification region of organoid on left), or (**B**) buffer with sodium (Na) +/- cytochalasin B (CB), or sodium-free buffer +/- cytochalasin B. Scale bars represent 200µm (**A** and **B** whole organoid images) and 100µm (higher magnification region in **A**). Bar chart (**C**) depicts quantification of green fluorescence within tubules of the representative experiment from (**B**) shown as a percentage of the total organoid area. Error bars represent standard error of the mean (SEM) from of n = 4 replicate organoids per condition. **D.** Schematic of the Transwell validation experiments using glucose-modified media. PT-EKO were generated and transitioned to 5mM or 30mM conditions before harvesting of PT-EKO tissues and corresponding culture media for transcriptional profiling and KIM-1 protein quantification, respectively. **E.** Soluble KIM-1 protein levels in 5mM (light yellow) and 30mM (dark yellow) glucose media collected from PT-EKO cultures, quantified using a solid-phase sandwich enzyme-linked immunosorbent assay (ELISA) kit. Data represent 4 independent experiments. Error bars indicate SEM of 4 measurements derived from 3 pooled biological replicates (note: n = 2 for the 5mM condition in Exp 3 due to a failed measurement). Statistical significance was assessed using a mixed-effects two-way ANOVA with Sidak multiple comparison test, comparing the mean of 5mM vs 30mM glucose within each experiment. **F.** Heatmap showing log-transformed transcripts per million for the top 20 differentially-expressed genes based on log fold change within PT-EKO exposed to 5mM or 30mM glucose-containing media (n = 3 – 4 biological replicates per condition across 2 independent replicate experiments, adjusted P value <0.01, mean normalized counts greater than 100).

To establish glucose transport specificity in PT-EKOs, the uptake of 2-NBDG was assessed in the presence and absence of sodium (required for optimal SGLT activity ^48^) and cytochalasin B (a GLUT inhibitor ^55,56^) (Figure 2B). Uptake of 2-NBDG was visibly reduced both in conditions lacking sodium (Na) and conditions exposed to cytochalasin B (CB) (Figure 2B). These results were validated by image quantification of 2-NBDG within tubules of the organoids as a proportion of total organoid area, confirming significant reduction in 2-NBDG fluorescence in both the absence of sodium and presence of 30µM cytochalasin B (Figure 2C). Despite low expression of SGLT genes and a lack of continuous apical / lumen-directed fluid flow, these results were suggestive of functional SGLT transporter proteins, with 2-NBDG uptake facilitated by the ability of such substrates to accumulate within tubule lumens via the ‘leaky’ or incompletely closed structures that occur in kidney organoids. Reduced 2-NBDG uptake in cytochalasin B conditions was supportive of the S3-active GLUT1 transporter, highly expressed within PT-EKO (Figure 1C). Despite GLUT1 being more often associated with basolateral glucose efflux *in vivo*, this transporter is also proposed to mediated glucose uptake from the peritubular space ^45^.

Given the known sensitivity of the PT to glucose levels (reviewed in ^57^), glucose responsiveness was explored by exposing PT-EKO to higher (30mM) and lower (5mM) glucose levels. PT-EKO were cultured in variable glucose conditions from D13+8-10 to facilitate PT elongation, with 2 days of agitation prior to harvest of tissue and culture media samples (Figure 2D). Imaging confirmed that proximalised elongated tubules were retained across all conditions (Supplementary Figure 2B). However, 30mM glucose conditions displayed altered morphology, with less distinct tubules, more frequent clusters of peripheral interstitial cells, and a darkened central core region (Supplementary Figure 2C). Previous studies have reported increased expression of KIM-1 / *HAVCR1*, a known clinical biomarker of kidney PT injury in humans ^58^, within kidney cells and organoid tissue following exposure to elevated glucose levels ^59,60^. To assess whether a similar response could be detected, an approach more akin to clinical KIM-1 assays was utilized, involving measuring shed KIM-1 protein ectodomain within culture media of organoids exposed to 5mM and 30mM glucose concentrations ^61^. In agreement with increased tissue expression of the *KIM-1* gene (*HAVCR1*) (Supplementary Figure 1D), KIM-1 solid-phase sandwich immunosorbent assay (KIM-1 ELISA) showed increased KIM-1 ectodomain in 30mM glucose conditions in three of four experiments (Figure 2E).

For experiments reaching significance, bulk RNA sequencing (bulk RNAseq) of organoid tissue samples was performed. Differential expression analysis identified significant increases a number of genes known to be involved in glucose and injury responses within the 30mM conditions of both experiments (Figure 2F). Examples included *TXNIP* and *ARRDC4* (insulin pathway suppressors involved in high glucose stress response, showing increased expression in cell models *in vitro* and rodents in response to elevated glucose ^62–65^), *CCL2* and *C2* (PT genes found to be elevated in diabetic patients and experimental animal models ^66–69^), and *BCL3* and *MX1* (associated with metabolic abnormalities or induced during kidney injury in tubular cells, with MX1 correlating with increased KIM-1 within AKI mouse models ^70–73^). Despite this glucose responsiveness, further analyses using curated injury markers from literature suggested experimental variability in the initiation of classic injury signatures, potentially arising from maturational differences or experiment duration and the ability to detect soluble KIM-1 ahead of some injury markers ^60^ (Supplementary Figure 2E).

### PT-EKO recapitulate metabolic characteristics of native PTs

Despite its ongoing role in glucose uptake, the maturing PT undergoes a metabolic shift away from glycolysis around the time of birth, becoming reliant on fatty acid oxidation and oxidative phosphorylation to fulfil its high energy demands ^74,75^. This is facilitated by large volumes of mitochondria within PT cells that exhibit dynamic spatial organization according to developmental stage, cortical vs medullary localization, and the onset of nephron filtration/flow ^76,77^. To explore mitochondrial abundance and dynamics in PT-EKO, organoids were examined for expression of TOMM20 (Translocase Of Outer Mitochondrial Membrane 20) following culture in standard static

Transwell (air-liquid-interface [ALI]) conditions, as well as after 48 hours of media flow (20µL/hour) within Ibidi 3D perfusion slides. Across both conditions, abundant TOMM20+ mitochondria were observed within HNF4A+/LTL+ PT cells (Figure 3AB, Supplementary Figure 3AB). PT-EKO cultured in standard static Transwell conditions exhibited dense mitochondria staining predominantly dispersed throughout the cytoplasm of PT (Figure 3A, Supplementary Figure 3A), with abundant mitochondria also present within distal tubules as previously reported ^77,78^ (Supplementary Figure 3C). When transferred to flow conditions, PT-EKO showed increased and highly-polarised EPCAM expression and frequently clustered mitochondria on the basolateral or apical membranes of HNF4A+/LTL+ PT (Figure 3B, Supplementary Figure 3B). Within flow conditions, mitochondrial localisation was found to be slightly more apical than basolateral, with quantification revealing higher levels of TOMM20+/LTL+ co-localisation compared to TOMM20+/EPCAM+ (LTL-) co-localisation (Figure 3C).

**Figure 3:**
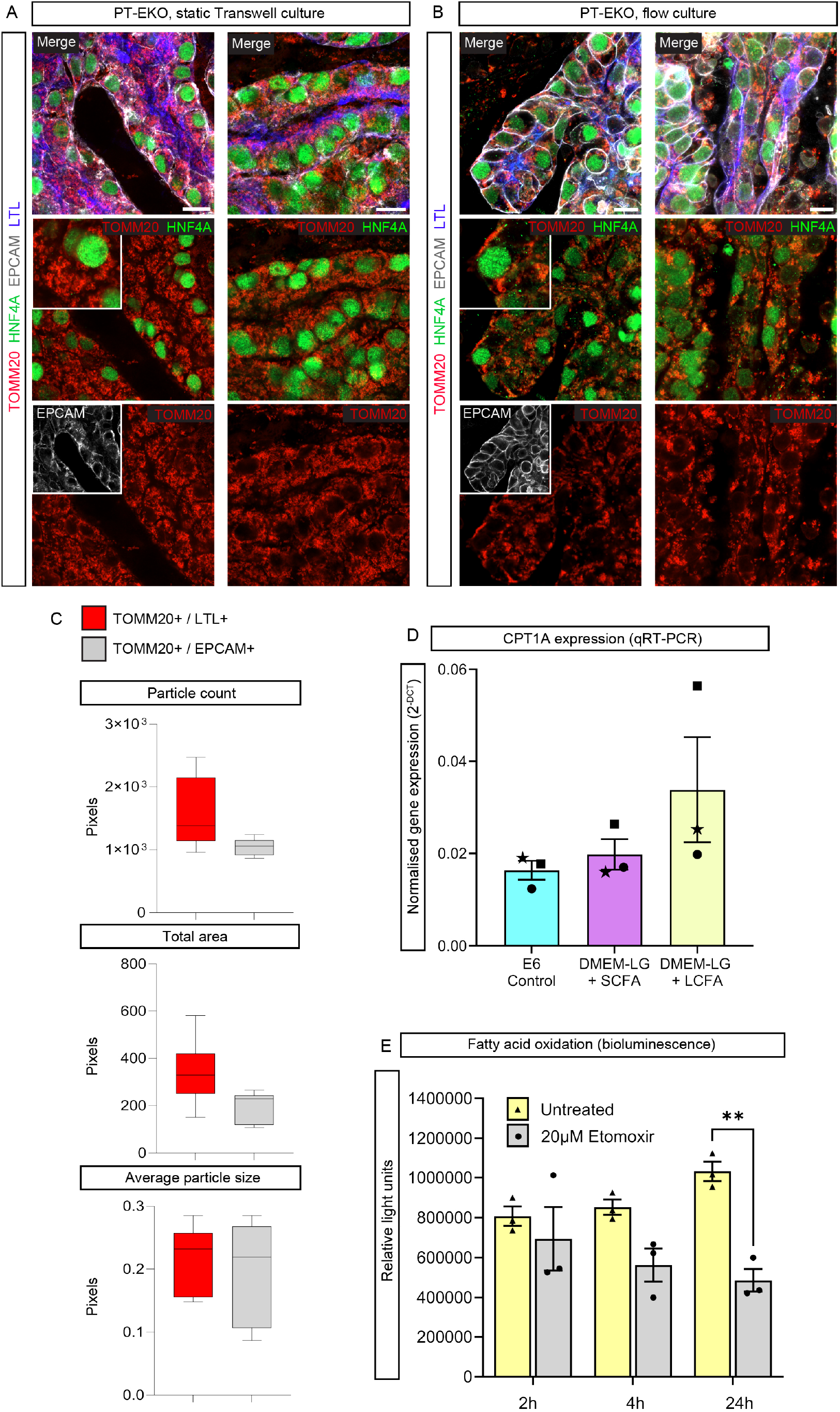
Metabolic characteristics of PT-EKO resemble those expected of native PT. **A-B.** Confocal immunofluorescence images of D13+18 PT-EKO cultured in standard static conditions on Transwell membranes (**A**) and under 20µL/hour media flow within Ibidi 3D Perfusion Slides (**B**) and stained with antibodies specific for Translocase of the Outer Mitochondrial Membrane 20 (TOMM20; red), proximal tubules (LTL; blue, Hepatocyte Nuclear Factor-4 alpha [HNF4A]; green), and nephron epithelium (EPCAM; grey). Insets depict regions of the same underlayer image at higher (middle row insets) or equal (bottom row insets) magnification. Scale bars represent 10µm. **C.** Quantification of TOMM20+ colocalization with LTL+ (red) and EPCAM+ (grey) particles, respectively. Data are depicted as pixels across particle count, total area, and average particle size plots. **D**. Quantitative real-time reverse transcription PCR (qRT-PCR) of D13+14 PT-EKO cultured in standard TeSR-E6 media (E6 Culture; cyan bar), DMEM-low glucose (LG) media supplemented with short chain fatty acids (SCFA), sodium butyrate and sodium acetate, (DMEM-LG + SCFA; magenta bar), and DMEM-low glucose (LG) media supplemented with long chain fatty acid (LCFA), palmitate, (DMEM-LG + LCFA; yellow bar). Error bars represent standard error of the mean (SEM) from n = 3 independent replicate experiments, identified by data point symbol, each datapoint representing the average of n = 3 biological replicates. **E.** Quantification of fatty acid oxidation (FAO) activity using a luciferase-based FAO reporting substrate within untreated (grey bars) and Etomoxir (ETO)-treated (yellow bars) PT-EKO cultured in DMEM-LG+LCFA. Error bars represent SEM from n = 3 organoids per condition across 3 separate experimental timepoints (2, 4, and 24 hours). Statistical significance was assessed using a two-way ANOVA with Sidak’ multiple comparisons test. Asterisks (**) denote P value ≤ 0.002 (P = 0.0017).

While abundant mitochondria are indicative of potential oxidative metabolic capacity, previous work has shown that PTs of standard kidney organoids are not capable of metabolizing long chain fatty acids (only short chain) due to a lack of carnitine palmitoyltransferase 1A (CPT1A); a rate-limiting mitochondrial outer-membrane enzyme critical for catalyzing long chain fatty acid shuttling into the mitochondria for oxidation ^75,79^. To assess the metabolic maturity of PT-EKO, PT clusters were assessed for their expression of fatty acid metabolism-related genes, including *CPT1A*, Acetyl-CoA acyltransferase 2 (*ACAA2*; rate-limiting enzyme [thiolase] catalysing final step of β-oxidation ^80,81^), diazepam-binding inhibitor / acyl-CoA binding protein (*DBI* / *ACBP* ; binds & transports long chain fatty acyl-CoA esters and is required for their availability for biological processes ^82,83^), *SLC27A2* (FATP2; apical PT transporter involved in long chain fatty acid transport into the cell ^84^), and *SLC16A1* (MCT1; PT membrane transporter involved in short chain fatty acid transport ^85,86^) (Supplementary Figure 3D). PT-EKO showed strong *CPT1A* expression in ∼20% of early PT populations (cluster 15, 1, and 3) and abundant expression of other fatty acid metabolism-related genes, *ACAA2, DBI, SLC27A2, and SLC16A1* (Supplementary Figure 3D).

These findings were strengthened by exposure of PT-EKO to altered metabolic substrates. PT-EKO were exposed to long- or short-chain fatty acids (LCFA or SCFA) for 4 days from days 8-10 of organoid culture and assessed for *CPT1A* expression. PT-EKOs exposed to LCFA consistently increased *CPT1A* (carnitine palmitoyltransferase 1A) expression across independent replicate experiments (Figure 3D). This increase was higher than SCFA-containing media, consistent with the role of CPT1A in the oxidation of LCFA and metabolic characteristics of *in vivo* PT. To establish whether *CPT1A* expression in lower glucose / LCFA conditions correlated with functional fatty acid oxidation (FAO), the activity of the pathway was measured using a luciferin-based assay (FAO-Glo Fatty Acid Oxidation Assay) that utilises a synthetic FAO substrate, capable of luminescence upon oxidation, and Etomoxir (ETO), an irreversible inhibitor of CPT1. FAO luminescent signal detected in untreated PT-EKO was found to be reduced following ETO treatment at all three timepoints, reaching statistical significance after 24 hours (Figure 3E). Together, these findings suggested that *CPT1A* gene expression correlated protein function and activity of the long chain FAO pathway within these PT-EKO.

### PT-EKOs demonstrate amenability to scale-up through progenitor cryopreservation

The viability of cell models for translational applications is advanced by the ability to bulk-up cell types and tissue of interest and cryopreserve until needed, facilitating shipment capacity for external manufacturing processes. To improve translation potential of our model, we tested scale-up of D13 monolayers and cryopreservation for later organoid generation. Cryopreservation of the monolayers was performed using commercial freezing solution (Cryo-SFM) compared to traditional cell cryopreservation methods using 10% dimethyl sulfoxide (DMSO). Following cryopreservation, thawed D13 monolayer cells were used to generate organoids for comparison back to controls (organoids generated from a subset of the same batch of monolayers that had not been frozen).

While viability of D13 cells at thaw was not significantly different between Cryo-SFM and DMSO conditions, the viability of D13 cells cryopreserved in DMSO was more variable between replicate experiments (Figure 4A). Brightfield and immunofluorescence imaging of nephron epithelium demonstrated characteristic aligned nephron spatial organization across all conditions and expression of standard patterning markers for distal and proximal nephron segments (Figure 4BC). However, compared to controls (generated with non-cryopreserved D13 monolayers), use of DMSO resulted in organoids with shorter tubules, larger NPHS1+ glomeruli, less SLC12A1+ loop of Henle (TAL), and changes to the central core region of the organoid shown previously to contain cortical stroma and pre- cartilage cells ^41^ (Figure 4C). In contrast, organoids generated from monolayers cryopreserved in Cryo-SFM showed higher similarity to non-cryopreserved controls with respect to morphology and patterning (Figure 4C).

**Figure 4:**
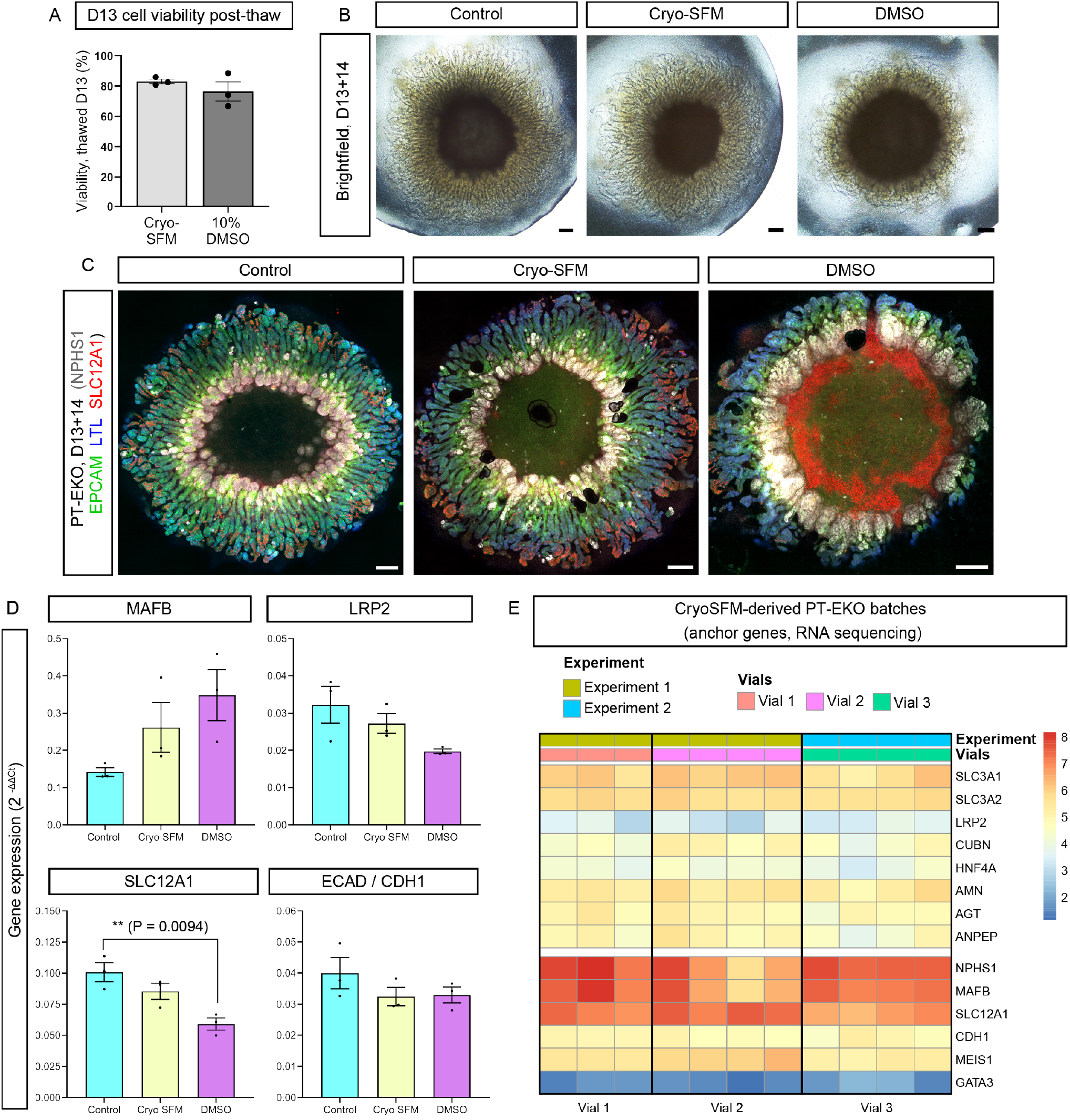
PT-EKO progenitors are amenable to cryopreservation at D13 and retain organoid differentiation capacity. **A.** Quantification of the viability of D13 organoid progenitors following thaw using the Countess™ 3 FL Automated Cell Counter, comparing D13 progenitors cryopreserved in Cryo-SFM (light grey bars) and 5% DMSO (dark grey bars). Error bars indicate SEM from 3 independent thawed D13 cryovials. **B-C**. Brightfield (**B**) and confocal immunofluorescence (**C**) images of PT-EKO at D13+14 generated from fresh D13 monolayers (control) or thawed D13 monolayers that had been cryopreserved in Cryo-SFM or DMSO. Immunofluorescence (**C**) depicts PT (LTL; blue), podocytes (NPHS1; grey), loop of Henle TAL (SLC12A1; red), and nephron epithelium (EPCAM; green). Scale bars represent 200µm. **D.** qRT-PCR of nephron patterning genes (*MAFB*; podocytes, *LRP2*; PT, *SLC12A1*; loop of Henle TAL, and *ECAD*; distal nephron) in D13+14 control PT-EKO (cyan bars), and PT-EKO generated from Cryo-SFM (yellow bars) and DMSO (magenta bars) cryopreserved D13 monolayers. Error bars indicate SEM from n = 3 biological replicates. Statistical significance was assessed using a one-way ANOVA with Tukey’s multiple comparisons test. Asterisks (**) denote two-tailed P value ≤ 0.01 for pairwise comparison. **E.** Heatmap showing expression of kidney anchor genes for PT (*SLC3A1/2, LRP2, CUBN, HNF4A, AMN, AGT, ANPEP*), loop of Henle TAL (*SLC12A1*), distal nephron (*ECAD, GATA3*), and stroma (*MEIS1*) within PT-EKO derived from 3 separate vials of cryopreserved monolayers across 2 independent experiments. Gene expression depicted as log-transformed transcripts per million.

These observations were confirmed by gene expression analyses (Figure 4D). Cryopreservation of D13 differentiated monolayers in Cryo-SFM was found to produce PT-EKO with gene expression levels more similar to non-cryopreserved controls compared to DMSO cryopreservation across three independent experiments (Figure 4D and Supplementary Fig. 4A). Increased *MAFB* expression (marking podocytes) and decreased *SLC12A1* expression (marking the loop of Henle TAL) were observed across multiple experiments and freezing conditions, indicating shifts in these cell populations (Figure 4D and Supplementary Fig 4A). However, *LRP2* expression in Cryo-SFM conditions remained similar to control levels across all three experiments, indicating retained proximal tubule proportions in these organoids and their amenability to scale-up through progenitor cryopreservation (Figure 4D and Supplementary Fig 4A).

To establish batch variability, bulk RNAseq analyses of PT-EKO derived from Cryo-SFM monolayers were performed. Analyses of PT-EKO derived from 3 separate vials of Cryo-SFM-preserved progenitors revealed similar expression levels of kidney anchor genes marking PT (*SLC3A1/2, LRP2, CUBN, HNF4A, AMN, AGT, ANPEP*), loop of Henle TAL (*SLC12A1*), distal nephron (*ECAD, GATA3*), and stroma (*MEIS1*) between vials (Figure 4E). Variability in *NPHS1* and *MAFB* expression was observed within replicates and between vials (Figure 4E), suggesting a level of inconsistency between podocyte proportion and/or development between replicates across batches, while tubule patterning showed minimal variability.

### Model versatility facilitates isolation of high-quality and expandable PT cells

2D PT cell cultures are often still preferred for translational applications such as drug screening owing to a desire for more simplistic, defined, and predictable cultures ^87^. However, they face challenges such as de-differentiation in culture, often associated with changes in substrate properties and loss of regulatory transcription factors such as HNF4A ^88^. Given our ability to generate scalable and functional PT-EKOs with distinct PT identities, we explored whether PT cells could be isolated and expanded, whilst retaining identity characteristics.

Consistent with our previous findings utilising the HNF4A:YFP reporter line ^41^, PT-EKO at D13+14 were found to contain approximately 20% PT cells (LTL+ / EPCAM+) (Figure 5A, Supplementary Table 2). Following FACS isolation, PT cells plated on collagen IV (plastic plates) or Geltrex (glass plates) adopted a cobblestone / cuboidal morphology characteristic of PT cells in 2D ^15,89^, highly similar to cultures of commercial PT lines (immortalised HK2 cells and primary human renal proximal tubule epithelial cells [hRPTECs]), and retained across multiple passages (Figure 5BC).

**Figure 5:**
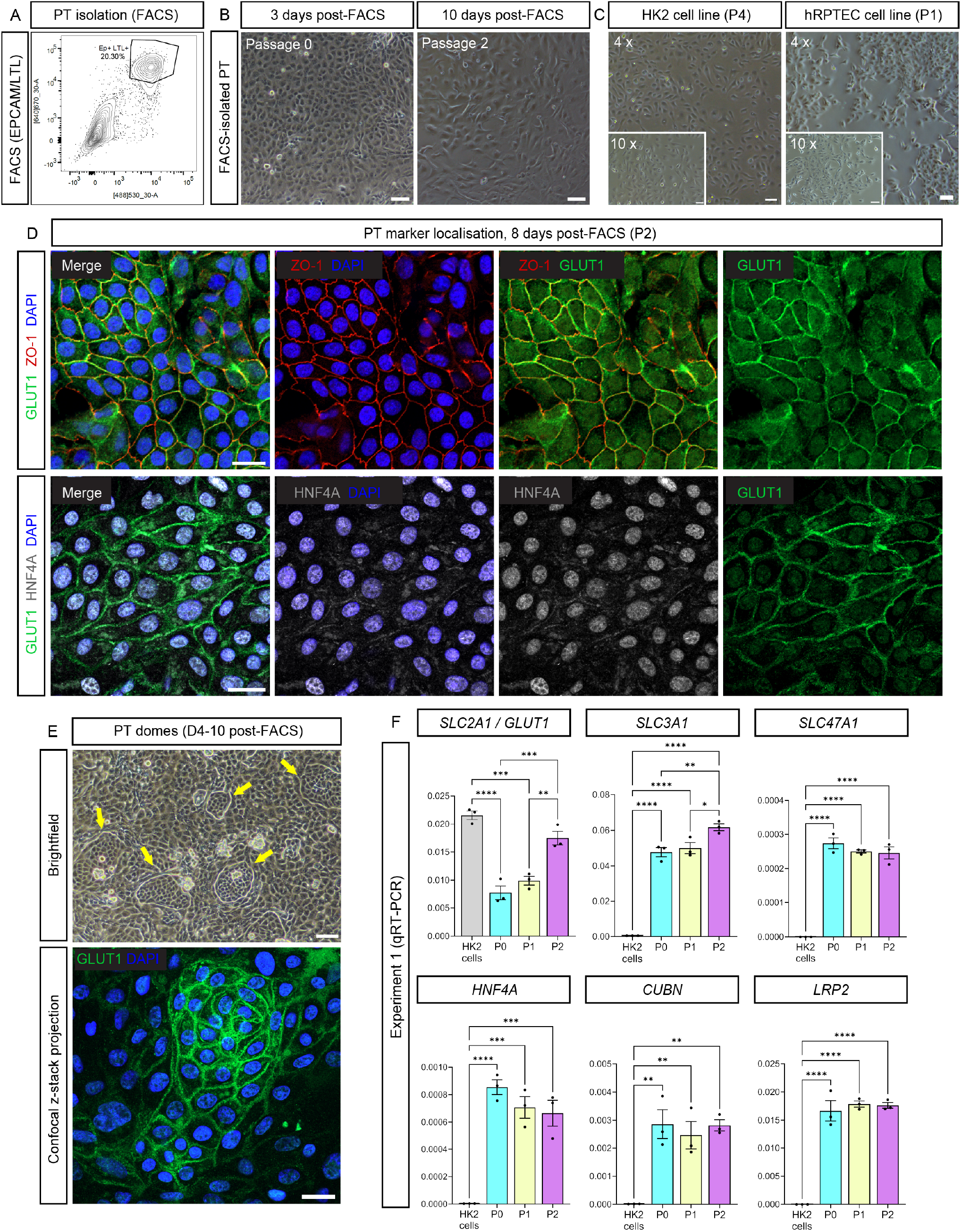
Isolated PT cells from D13+14 PT-EKOs retain PT characteristics across multiple passages. **A.** FACS plot example showing the isolated LTL+/EPCAM+ PT population derived from D13+14 dissociated PT-EKO prior to re-culture. **B**. Brightfield images of PT cells isolated from PT-EKO at 3 days (passage 0 [P0]) and 10 days (passage 2 [P2]) days post-FACS. Scale bars represent 100µm. **C.** Brightfield image of commercially-available immortalised (HK2; left) and primary (hRPTEC; right) cell line cultures. Insets depict cells at higher magnification. Scale bar represents 100µm. **D**. Confocal immunofluorescence images of isolated PT cultures at 8 days post-FACS (passage 2 [P2]) depicting tight junction protein expression (ZO-1; red), membrane-bound glucose transporter expression (GLUT1; green), nuclear transcription factor HNF4A (grey), and cell nuclei (DAPI; blue). Scale bars represent 20µm. **E**. Brightfield (top) and confocal immunofluorescence (bottom) images depicting formation of domed structures in PT cultures at 4 – 10 days post-FACS. Domes indicated with yellow arrows in top brightfield image express GLUT1 protein (bottom image; green). Scale bars represent 100 µm (top image) and 20 µm (bottom image). **F**. qRT-PCR of PT genes in isolated PT cells across a 7-day time course (passage 0 [P0; cyan bars], passage 1 [P1; yellow bars], passage 2 [P2; magenta bars]) compared to HK2 cells (grey bars). Error bars represent SEM of n = 3 biological replicates. Statistical significance was assessed using a one-way ANOVA with Tukey’s multiple comparisons test. Asterisks denote two-tailed P values of ≤0.05 (*), ≤0.01 (**), ≤0.0001 (***), and ≤ 0.0001 (****) for pairwise comparisons.

Immunofluorescence of the isolated PT cells at passages 0, 1, and 2 (0 – 10 days post-FACS) demonstrated retained expression and expected cellular localisation of multiple PT markers (Figure 5D and Supplementary Figure 5A). This included distinct membrane expression of ZO1 (tight junction protein critical for apical-basolateral separation and epithelial barrier function ^90^) and the basolateral membrane glucose transporter, GLUT1. Importantly, bright nuclear expression of HNF4A, required for cultured PT identity, was also maintained at all timepoints (Figure 5D, Supplementary Figure 5A). Upon reaching confluence, PT cultures were also found to produce and retain numerous raised dome structures with high levels of membrane-bound GLUT1 transporter, indicative of active transepithelial transport (Figure 5E).

Initial assessments of PT gene expression levels via qRT-PCR further strengthened the PT-like identity of the isolated cells (Figure 5F, Supplementary Figure 5B). Functionally-critical PT markers across 3 timepoints post-FACS were assessed, including *SLC2A1* (encoding the GLUT1 glucose transporter ^45,91^), *SLC3A1* (encoding the rBat cystine and di-basic amino acid transporter responsible for type I cystinuria ^92^), *SLC47A1* (encoding cationic drug and toxin extrusion protein, MATE-1^93^), *HNF4A* (encoding the HNF4A transcription factor critical for retained identity of PT cells in culture and their development *in vivo* ^88^), and *CUBN* / *LRP2* (encoding the CUBILIN-MEGALIN [CUBN-MEG] transporter complex critical for preventing urinary protein loss ^94^) (Figure 5F). Gene expression levels were either maintained or increased in the isolated PT cells across the 10 day duration of the experiment and multiple dissociation / passaging steps, with a consistent increase in glucose and amino acid transporters, *SLC2A1* and *SLC3A1*, as well as higher *SLC3A1, SLC47A1, HNF4A, CUBN* and *LRP2* in isolated PT cells compared to immortalized HK2 cells (Figure 5F, Supplementary Figure 5B).

These characteristics were found to be partially retained in thawed and re-cultured PT-EKO-derived PT cells following cryopreservation in Cryo-SFM (Supplementary Figure 6A-E). At 3 days post-thaw, cryopreserved PT cells regained a characteristic cuboidal morphology and showed dome formation indicative of transepithelial transport (Supplementary Figure 6AB). While expression of some PT genes (*CUBN, LRP2, HNF4A, SLC3A1* and *SLC47A1*) were decreased in PT cells post-thaw, expression levels remained higher than immortalized HK2 cells and cultures showed similar expression of ZO-1, GLUT1, and HNF4A proteins via immunofluorescence to freshly isolated PT cells (Supplementary Figure 6C-E).

To more comprehensively investigate PT identity and transporter activity, isolated PT cells were assessed with substrate uptake assays and further marker analyses, benchmarking against commercially-available primary human renal epithelial cells (hRPTECs; isolated from human adult kidney cortex) of similar passage numbers (Figure 6A-D). Isolated PT cells were found to retain robust uptake of three fluorescence substrates after 3 passages, including TRITC-conjugated albumin (transported by the MEG-CUBN complex ^41^), 4–4-dimethylaminostyryl-*N*-methylpyridinium (ASP; an organic cation surrogate, predominantly transported by OCT2 / *SLC22A2* ^95^), and 4′,6-diamidino-2-phenylindole (DAPI; an organic cation surrogate effectively transported in PT by the SLC47 family Multidrug and Toxin Extrusion proteins 1 [MATE-1 / *SLC47A1*] and 2-K [MATE2-K / *SLC47A2*] ^96^) (Figure 6A). Confirming the specificity of uptake, both ASP and DAPI uptake were effectively inhibited or reduced by treatments with the organic cation transporter inhibitor, Cimetidine (Figure 6B). Transport of ASP and DAPI was only slightly higher in primary hRPTECs of a similar passage number compared to PT-EKO-derived isolated PT cells, while TRITC-albumin uptake was significantly increased in isolated PT cells, suggesting higher activity of the CUBN-MEG complex (Figure 6B).

**Figure 6:**
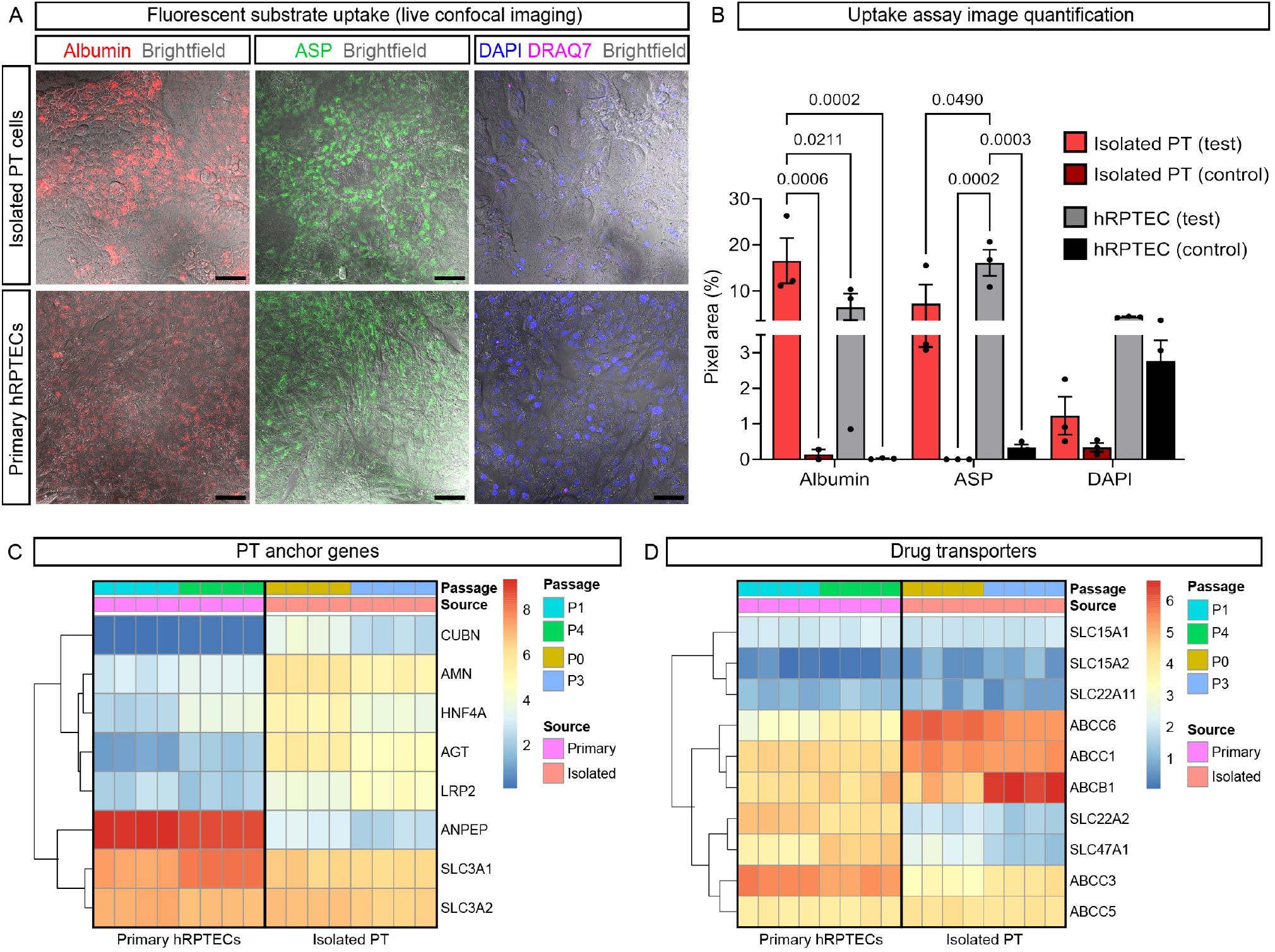
PT cells isolated from PT-EKO possess functional PT transporter expression. **A.** Live confocal images of isolated PT cells (top row) and commercially-available primary hRPTECs (bottom row) depicting uptake of TRITC-albumin (red, left images), ASP (green, middle images), and DAPI (blue, right images). Dead cells in DAPI image are labelled with DRAQ7 (magenta), confirming live cell uptake. All panels show brightfield overlay. Scale bars represent 50 µm. **B**. Quantification of segmented fluorescent signal in substrate uptake assay experiment shown in (**A**) across test (substrate only) and control (pre-incubated with inhibitor, Cimetidine, prior to substrate exposure) conditions. Data are depicted as fluorescent pixel area for n = 3 replicate wells per condition. Error bars represent SEM. Statistical significance was assessed using a two-way ANOVA with Sidak’s multiple comparison test. Numerical P values are shown above significant comparisons. **CD.** Heatmaps showing the expression of PT anchor genes (**C**) and PT drug transporters (**D**) in primary hRPTECs and isolated PT cells across 2 passages with n = 4 biological replicate wells per condition. Gene expression depicted as log-transformed transcripts per million.

Functional assay results were supported by transcriptional analyses of PT anchor genes. Although slight reductions in expression levels of some genes across passages was noted, such as *HNF4A* and *ANPEP* in hRPTECs and isolated PT cells (Figure 6C), isolated PT cells demonstrated higher expression of *CUBN* and *LRP2* overall (conferring the CUBN-MEG receptor complex for protein transport), as well as key genes responsible for PT transporter activities and retained PT identity *in vitro.* These included genes conferring proteins required for the regulation of sodium availability (*AGT*), membrane anchoring and receptor-mediated endocytosis (*AMN*), and transcriptional regulation of transport activities (*HNF4A*) ^97–101^ (Figure 6C). Higher expression was also noted for a number of PT drug transporters, particularly that of the ATP-binding Cassette (ABC) family of drug transporters which play a critical roles in transporting a very broad range of drug substrates and conferring drug resistance, including *ABCB1* (*Multiple Drug Resistance 1* [*MRP1*] / P-glycoprotein [P-gp]) and *ABCC1* (Multidrug Resistance Protein 1 [MRP1]) ^102,103^ (Figure 6D). While lower expression of organic cation transporter genes *SLC22A2* (OCT2) and *SLC47A1* (MATE-1) were observed in isolated PT cells compared to primary hRPTECs, isolated PT cells showed robust and specific uptake of their respective transporter substrates, ASP and DAPI (Figure 6A), suggesting a level of functional expression.

Finally, given the more S3-like identity of the PTs within PT-EKO, isolated PT cells from these organoids were analysed for their expression of PT segment-specific gene signatures via bulk RNAseq across 2 passages, comparing to hRPTECs (Supplementary Figure 6F). Expression of a broad range of earlier (S1-S2) and later (S2-S3) PT markers were assessed, curated from mammalian gene and protein distribution studies, with a preference for human studies where available (Supplementary Table 1). Compared to primary hRPTECs, gene expression analyses of isolated PT indicated higher expression levels of PT markers expected to be enriched towards the later S2-S3 segments of PTs *in vivo* (Supplementary Figure 6F), suggesting that isolated PT cells retained a more S3-like identity following derivation from PT-EKO.

## DISCUSSION

The optimal replication of PT function, injury responses, and disease mechanisms *in vitro* depends on the development of authentic PT models that accurately reflect human physiology. Achieving this has remained challenging, largely due to the exceptional structural, metabolic, and functional complexity of the human PT, which exhibits marked cellular heterogeneity and segment-specific specialisation along its length. In this study, we demonstrate that PT-EKOs are enriched for a functionally-critical PT cell population that is most prone to injury; the S3 segment. This enrichment is accompanied by transcriptional alignment with adult human PT profiles and evidence of physical PT segmentation, highlighting the uniqueness of this *in vitro* model.

Recapitulation of key human PT features was further supported by appropriate transport and metabolic activity, as well as responsiveness to environmental cues. PT-EKOs demonstrated the capacity to respond to long-chain fatty acids through upregulation of *CPT1A*, the rate-limiting enzyme in mitochondrial long-chain fatty acid oxidation, coupled with fatty acid oxidation pathway activity. This response was consistent with the known metabolic profile of maturing PTs *in vivo*, which rely heavily on fatty acid oxidation to meet their substantial energy demands (reviewed in ^76,104^) and was strengthened by their high levels of TOMM20+ mitochondria. Consistent with less mature PTs residing in deeper cortical zones of developing mouse and human kidneys ^77^, PT-EKOs cultured on Transwell membranes at an ALI exhibited dispersed mitochondrial localisation throughout PT cell cytoplasm, as well as abundant mitochondria in distal PT-EKO tubules as observed *in vivo* ^78^. As PTs mature, the onset of fluid flow has been suggested to induce apical-to-basolateral mitochondrial re-distribution ^77^. Although not strictly basolaterally positioned as reported in actively filtering PTs in developing mouse and human ^77^, PT-EKO cultured in OoC platforms developed a combination of apical and basolateral mitochondria, potentially reflecting early changes in PT maturity. Whether the basolateral restriction of mitochondria could be enhanced through the application of a more direct luminal flow, along with refined flow rates, flow duration, and metabolic substrate exposure, warrants further investigation.

Given the PTs vulnerability to injury and disease, the creation of accurate PT models remains a high-priority application of kidney *in vitro* systems. The development of S3-enriched platforms would be particularly advantageous, given the heightened susceptibility of this segment to ischemic and hypoxic injury, drug-induced nephrotoxicity (reviewed in^6^), and metabolic disturbances, including fatty acid and glucose levels. Of relevance, recent analyses have identified a higher hypoxia-associated signature in PT-EKOs compared to some alternate organoid protocols ^105^. While identification of distinct PT subpopulations may have been challenged in this case by reduced resolution and a broader sub-setting approach, when combined with the findings of the current study, it raises the possibility S3-like PT cell enrichment may contribute to this susceptibility.

Furthermore, the S3 segment has also been associated with early molecular changes that may precede and predict later pathophysiological changes in diseases such as diabetes ^7^. While hyperglycaemia is a major driver of AKI and CDK globally, relatively few studies have examined the ability of kidney organoids to model hyperglycaemic injury ^106,107^, nor examined correlations with clinically-relevant soluble urinary/serum biomarkers. In exploring the utility of PT-EKO, the current study demonstrated the functionality and specificity of glucose transporters within this model. Despite a level of variability in injury signatures between experiments, in agreement with soluble KIM-1 elevations observed in diabetic patient serum/urine ^108,109^, responses of high-glucose–exposed kidney organoids reported by Garreta et al. ^106^, and observations from diabetic mouse PTs ^110,111^, PT-EKOs showed appropriate responses to elevated glucose concentrations, including upregulation of glucose-responsive genes and detectable KIM-1 ectodomain within culture media samples. Interestingly, PT-EKO also possessed higher *GLUT1* expression within S3-like PT populations compared to adult human kidneys. This may have resulted environmental factors, such as higher-than-physiological glucose levels within standard culture media or hypoxia influencing *GLUT1* elevation in the S3-like PT cell population (confirmed to also express non-hypoxia-associated S3 genes including ALPI ^112^). Together, these findings demonstrate the utility of the model in capturing early injury responses, while highlighting that further recapitulation of the in vivo kidney environment may improve the reproducibility of injury signatures.

Despite the anticipated benefits of increasingly complex 3D kidney structures for more physiologically relevant modelling, homogeneous PT cell populations are still seen as advantageous considering their compatibility with quality control, screening, expansion, and storage methodology ^87^. Given the limitations of primary and immortalised PT cell lines, rapid advances in stem cell technologies have initiated multiple developments in alternative, scalable sources of PT cells, including direct differentiation from ESCs or iPSCs ^19,20,113^ and isolation of PT cells from PSC-derived kidney organoids ^26,31,37,114^, with such approaches facilitating renewable and genetically-defined cell sources able to be further engineered and expanded to PT identities. However, a challenge remains in demonstrating retained PT characteristics across serial passaging and cryopreservation, including correct morphology, growth characteristics, and expression of PT-instructive transcription factors. In the current study, the translational utility of PT-EKOs was expanded by their capacity to serve as an enriched source of isolatable high-quality PT cells, comparable to or improved beyond their commercial counterparts across a number of PT cell characteristics, yet showing a more S3-like gene signature. PT cells isolated from PT-EKO could be expanded across multiple passages, retaining a number of critical PT-specific anchor genes, such as important transporter co-factors and functionality-driving transcription factors including *HNF4A* ^115^. Isolated PT cells also showed robust functionality of several transporters relevant to therapeutic development, including OCT2, MATE1, and the MEGALIN-CUBILIN complex, likely facilitated by the retained expression *AMN*, required for membrane anchoring of the complex ^97–99^. Given the promise of these early-stage investigations, further optimisations to improve the purity of this unique isolated PT cell population and determine the ideal growth conditions is likely to facilitate further improvements in transporter stability throughout passaging and post-cryopreservation.

Looking forward, the ability to model architecturally accurate 3D proximal nephrons with distributed functionally distinct PT subtypes remains an ultimate objective. Recent advances have enabled the generation of PT-enriched organoid models ^26,27,41^, with some evidence of nephron spatial organisation ^41^. However, the reproducible generation of the higher-order PT structural features required to maximise functionality remains unresolved. Bioengineering approaches such as 3D bioprinting of commercial PT cells within gelatin–fibrinogen matrices have opened up the possibility of directly engineering geometries like PT convolutions, also demonstrating the benefits of open, perfusable luminal architectures ^32^. Whether such architectural features can be achieved within self-organising stem cell–derived tissues will likely depend on improved understanding of nephron morphogenesis, including the interplay between morphogen gradients, mechanical forces, and cell–cell interactions thought to influence these physical nephron transitions ^116–118^.

In summary, here we present PT-EKOs as a unique PT model, showing S3-like cell enrichment and capturing structural, metabolic, and injury-responsiveness features of the human PT. The emergence of structurally subdivided PTs represents a distinct advance in modelling PT biology. Combined with serving as an expandable source of high-quality isolated PT cells, this platform provides a framework for interrogating segment-specific function and injury responses, holding promise for drug development, improving safety assessment, modelling kidney disease, and supporting future artificial kidney technologies of the future.

## EXPERIMENTAL PROCEDURES / METHODS

### iPSC maintenance & generation of PT-EKOs

Expansion and maintenance of the CRL1502.2 iPSC line (derived from WS1 CRL-1502 female fibroblasts [ATCC] using episomal reprogramming methods described in reference ^41^ [registration: MCRIi019-A, https://hpscreg.eu/cell-line/MCRIi019-A]) was performed as described previously ^43^. Briefly, cells were grown in Essential 8 Medium or Essential 8 Flex Medium (Thermo Fisher Scientific, Waltham, MA) on Matrigel- (Corning, NY, USA, cat # 354277) coated plates, passaging every 2-3 days with 0.5mM EDTA in 1X phosphate-buffered saline (PBS). For PT-enhanced kidney organoid generation, iPSCs were seeded into laminin-521 (BioLamina, Stockholm, Sweden, Cat# LN521) coated 12-well plates at a density of 30,000 cells/well and differentiated as a monolayer for 13 days (including 5 days of CDBLY2 exposure) prior to manual organoid generation according to methods detailed in reference ^41,43^.

### Single cell-RNA sequencing

ScRNAseq analyses of S1-S2 and S3 PT markers in PT-EKO were performed on our existing published single cell data set (GEO accession: GSE184928) ^41^. This data set consisted of >11,000 cells from four hashtag oligo barcoded replicates. Library demultiplexing in CellRanger, normalization, and marker analysis in Seurat (3.1.4) were performed as described previously ^41^.

For comparisons of S1, S2, and S3 proportions, the PT-EKO scRNAseq dataset above was compared to two standard Takasato protocol-derived kidney organoid scRNAseq datasets (D7+12 ‘D18 Howden’ GSE119561 ^50^ and D7+18 ‘D25 Wilson’ GSE161255 ^51^ ) and a D27 proximal-biased kidney organoid single nuclear RNAseq dataset (D7+20 ‘D27 Schnell’ [Day27Ly29], GSE290931) ^27^. PT and early PT cells within the datasets were scored for their expression of S1 (*SLC6A19, SLC36A2, RBP4, SLC2A2, SLC5A2, SLC34A1*), S2 (*CA4, LYZ, SLC6A20, SLC15A2*) and S3 (*SLC5A1, ALPI, SLC2A1, ALC6A20, SLC15A2*) marker genes using the UCell package ^119,120^. Cells were classed as positive with a score above 0.

For Pearsons correlation of PT-EKO PT populations with that of 2 adult human kidney reference datasets (Liao *et al.* [2020] and Lake *et al.* [2023, Kidney Cell Atlas] ^52,53^), PT-EKO PT populations showing similar identities based on PT segment-specific anchor gene expression (Supplementary Table 1), were merged to create S3-like and S1-S2 like populations (‘PTEKO_PT-A’ [merge of most similar S3-like clusters; cluster 3 and 8] and ‘PTEKO_PT-B’ / ‘PTEKO_PT-C’ [merge of most similar S1-S2-like clusters; clusters 1 and 13 (C), cluster 15 (B)]). To compare PT-EKO with adult PT populations, adult human datasets located at GSE131685 and https://www.kidneycellatlas.org/ were processed following the same parameters of the original publications ^52,53^. Correlation analysis was performed by sub-setting the PT clusters from all datasets, summing gene counts for all cells per cluster, and then calculating gene counts per million. The pseudobulk counts were then compared for each cluster using Pearson’s correlation.

Code for the scRNAseq analyses in the current study is available on request and will be made publicly available following publication through the Kidney Regeneration Github repository (https://github.com/KidneyRegeneration/Vanslambrouck2026).

### Bulk RNA sequencing

Bulk RNA expression libraries were generated using the Illumina stranded mRNA kit and sequenced on an Illumina Novoseq 6000 sequencer by the Australian Genome Research Facility (AGRF). Raw fastq files were mapped to human genome version GRCh38 using the nf-core/rnaseq pipeline version 3.12.0 (https://nf-co.re/rnaseq/3.12.0/), outputting tables of raw counts and gene transcripts per million. Differential expression analysis of high vs low glucose treatment was conducted on count data using DESeq2 (v1.49.5), including batch as a covariate in the design formula. Transcripts per million were log-transformed and used to plot heatmaps, using the pheatmap library (v1.0.13), with for filtered differential gene expression lists or curated gene lists.

### Immunofluorescence

Organoids underwent fixation and wholemount immunofluorescence as described previously ^43^ using the antibodies detailed in Table 1, diluted in 0.3% TX-100/PBS. Imaging was performed on the ZEISS LSM 900 confocal microscope (Carl Zeiss, Oberkochen, Germany) with acquisition and processing performed using ZEISS ZEN software (ZEN 3.4 Blue Edition [Zeiss Microscopy, Thornwood, NY]) and Fiji ImageJ (https://imagej.net/software/fiji/).

**Table 1:** Antibodies used for immunofluorescence.

| Specificity | Host species | Dilution range | Manufacturer and identifier |
| --- | --- | --- | --- |
| DAPI |  | 1:1000 | Thermo Fisher Scientific (D1306) |
| ECADHERIN | Mouse monoclonal IgG2a, Kappa | 1:300 | BD Biosciences (610181) clone 36 |
| EpCAM (Alexa488 or Alexa647 conjugate) | Mouse monoclonal IgG2a, Kappa | 1:300 | BioLegend (324210 and 324212) clone 9C4, Lot B352438 and B314235 |
| GATA3 | Goat polyclonal IgG or rabbit monoclonal IgG | 1:300 | R&D Systems (AF2605) and Cell Signalling Technology (5852S clone D13C9, Lot UZQ0219011) |
| GLUT1 | Mouse monoclonal IgG1kappa | 1:300 | Santa Cruz (sc-377228), Lot H1823 |
| HNF4A | Mouse monoclonal IgG2a or rabbit monoclonal IgG | 1:300 | Invitrogen (MA1-199) clone K9218, Lot WI3366551 and Cell Signalling Technology (3113) Clone C11F12 |
| NEPHRIN | Sheep polyclonal IgG | 1:300 | R&D Systems (AF4269), Lot ZMU0221031 |
| Proximal tubule brush border membrane | *Lotus tetragonolobus* lectin (LTL), biotinylated | 1:300 – 1:500 | Vector Laboratories (B-1325-2) |
| SLC12A1 | Rabbit polyclonal IgG | 1:300 – 1:400 | Proteintech (18970-1-AP), Lot 00045810 |
| TOMM20 | Rabbit monoclonal IgG | 1:300 | Abcam (ab186735), clone EPR15581-54, Lot 1097681-20 |
| ZO-1 | Goat polyclonal IgG | 1:300 | Invitrogen (PA5-19090), Lot UC283843 |
| Streptavidin, Alexa Fluor 405 conjugate | Biotin | 1:500 | Thermo Fisher S32351 |
| Alexa Fluor 488 Donkey anti-mouse IgG (H+L) | Mouse IgG (H+L) | 1:500 | Thermo Fisher A21202, Lot 2563848 |
| Alexa Fluor 568 Donkey Anti-Rabbit IgG Antibody | Rabbit IgG | 1:500 | Thermo Fisher A10042, Lot 2540901 |
| Alexa Fluor 568 Donkey anti-goat IgG antibody | Goat IgG (H+L) | 1:500 | Thermo Fisher A175704 |
| Streptavidin, Alexa Fluor 647 conjugate | Biotin | 1:300 | Thermo Fisher S21374 |
| Alexa Fluor 647 donkey anti-goat IgG (H+L) | Goat IgG (H+L) | 1:500 | Thermo Fisher A21447, Lot 2841625 |
| Alexa Fluor 647 Donkey anti-sheep IgG (H+L) | Sheep IgG | 1:500 | Thermo Fisher A21448, Lot 2563882 |

### Real-time quantitative reverse transcription PCR

RNA extraction, cDNA synthesis, and quantitative RT-PCR (qRT-PCR) were performed using the Bioline Isolate II Mini/Micro RNA Extraction Kit (Bioline, Meridian Bioscience, London, UK,, cat# BIO-52075), SensiFAST cDNA Synthesis Kit (Bioline, cat# BIO-6504), and the SensiFAST SYBR Lo-ROX Kit (Bioline, cat# BIO-94020) according to the manufacturer’s instructions. Triplicate technical replicates were performed for each reaction using the primer pairs detailed in Table 2. RT-PCR data were collected using QuantStudio^TM^ Design & Analysis Software (version 1.4.1) installed on the QuantStudio^TM^ 5 Real Time PCR System (ThermoFisher Scientific, MA, USA). Data were analysed using the delta-CT method, normalized to the expression of housekeeping gene *GAPDH*, and graphed and analyzed in Graphpad Prism version 10.1.2.

**Table 2:**
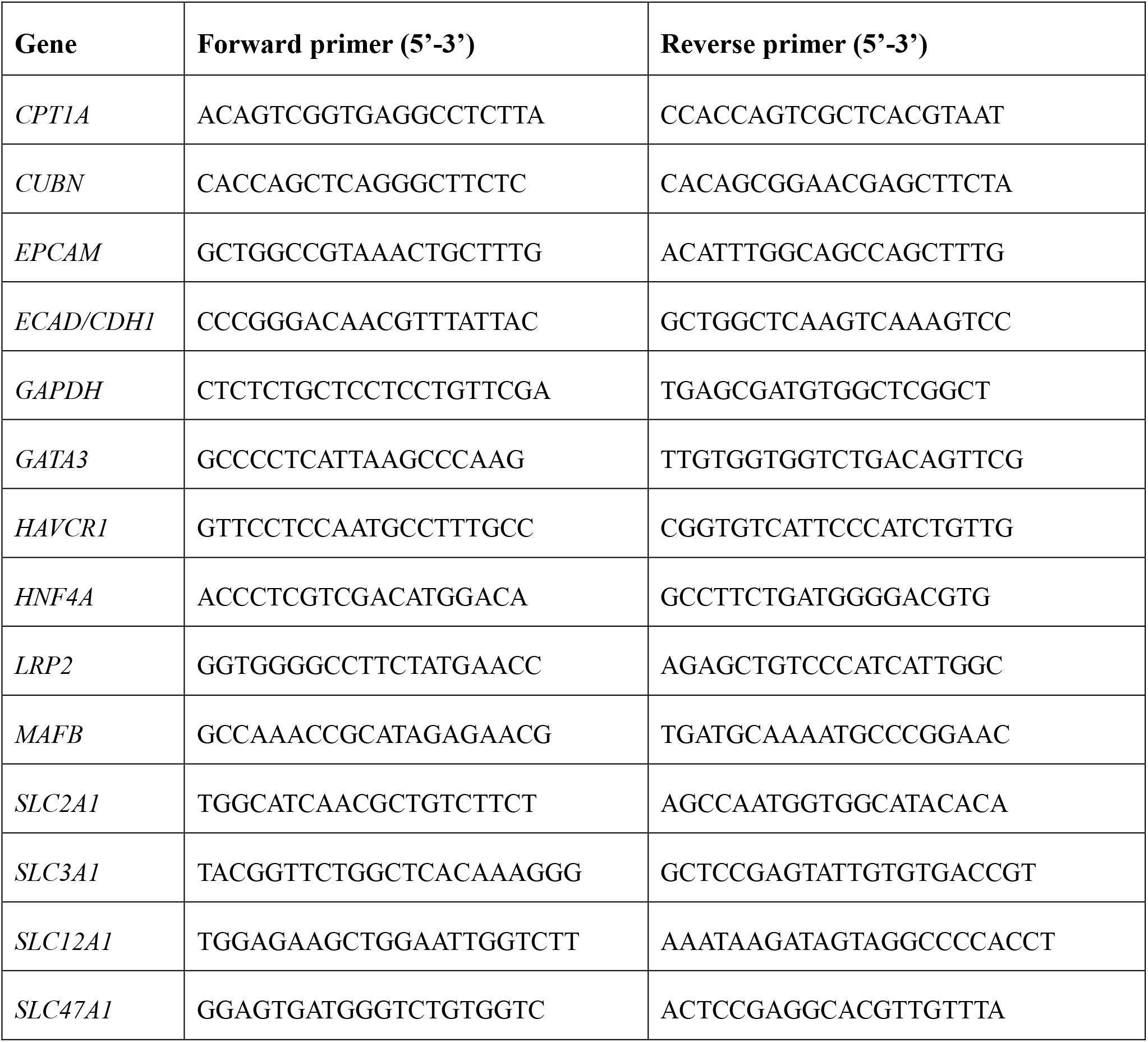
Primer sequences used for qRT-PCR.

### Glucose-modified culture conditions

To induce hyperglycemic injury, D13+8 - Day 13+10 PT-EKO were incubated in a 1:1 mixture of standard organoid culture media (TeSR-E6) and glucose-modified media (5mM or 30mM, detailed below) for 2 days, before refreshing with 100% glucose-modified media (5mM or 30mM). Glucose-modified media consisted of glucose-free and glutamine-free DMEM (Gibco, ThermoFisher, cat# A1443001) supplemented with 2.5 mM GlutaMAX (Gibco, ThermoFisher, cat# 35050061), 1% penicillin/streptomycin, and either 5mM or 30mM glucose (Sigma Aldrich, Merch, MA, USA, cat# G7528). PT-EKO were cultured in glucose-modified media for 6-8 days, refreshing every second day, prior to harvesting.

### Glucose uptake assay and quantification

Glucose transport activity was assessed following methods published previously ^54^. Briefly, D13+14-15 PT-EKO were incubated for 2 hours at 37°C in sodium buffer (140 mM NaCl, 5 mM KCl, 2.5 mM CaCl_2_, 1 mM MgSO_4_, 1 mM KH_2_PO_4_, and 10 mM HEPES [pH 7.4]) or sodium-free buffer (140 mM *N*-Methyl-d-glucamine, 5 mM KCl, 2.5 mM CaCl_2_, 1 mM MgSO_4_, 1 mM KH_2_PO_4_, and 10 mM HEPES [pH 7.4]) containing 200 µM of fluorescent D-glucose analogue (2-(N-(7-Nitrobenz-2-oxa-1,3-diazol-4-yl)Amino)-2-Deoxyglucose [2-NBDG], ThermoFisher Scientific, USA, cat# N13195). After incubation, organoids were washed 3 times (5 minutes each) in sodium-free buffer before transferring to glass-bottom plates for live imaging (ZEISS LSM 780 confocal microscope, 488-nm excitation and 545-nm emission).

Control conditions included PT-EKO without incubation in 2-NBDG (buffer incubation only) and organoids exposed to glucose uptake suppressors, including sodium-free buffer and 30µM cytochalasin B (Cayman Chemical, USA, cat# 11328). For uptake inhibition conditions, PT-EKO were incubated in either sodium-free buffer +/- cytochalasin B or sodium-containing buffer +/- cytochalasin B for 30 minutes prior to spiking with 2-NBDG and a further 2 hours incubation in standard cell culture conditions at 37°C). Following incubation, organoids were washed 3 times (5 minutes each) in their respective buffers before confocal live-imaging under the same 488nm excitation parameters as above. For quantification, live confocal images above were analysed using Fiji ImageJ 1.54p Labkit plug-in. A separate classifier was trained for the detection of 2-NBDG and applied to the 488 channel to distinguish positive signal from background. A custom macro was developed to batch analyse samples across multiple experiments. The area of the whole organoid was manually traced using the Freehand selection tool and total organoid area was quantified according to scale. The area occupied by the fluorescent tubules was quantified and expressed as a percentage of the total organoid area (defined manually) within each image.

Statistical analyses were performed using a two-way ANOVA (ordinary, ungrouped) followed by Sidak’s multiple comparisons post hoc test. Data are presented as mean ± SEM and differences considered statistically significant at P <0.05.

### ELISA quantification of KIM-1 protein

For quantification of soluble KIM-1, frozen culture media samples were thawed on ice and analysed using a KIM-1 solid-phase sandwich enzyme-linked immunosorbent assay (ELISA) kit following the manufacturer’s protocol (Thermo Fisher cat# EHHAVCR1). ELISA readings were obtained at 450nm absorbance using the Tecan i-Control software connected to Infinite 200 Pro plate reader (Tecan, Männedorf, Switzerland). A standard curve was fitted using a four-parameter algorithm as recommended by manufacturer, using GraphPad Prism’s Sigmoidal 4PL analyses function (GraphPad version 10.1.2).

### PT-EKO organ-on-a-chip (OoC) culture using perfusion slides

To compare flow and static conditions, D13+12 PT-EKO from the same D13 monolayer differentiation were cultured either on Transwell membranes (as per the PT-EKO generation protocol detailed above) for 48 hours without media change, or under flow conditions for an equivalent 48 hours at a rate of 20µL/hour. For flow-induction, the chambers of μ-Slide III 3D Perfusion chips (Ibidi, Gräfelfing, Germany, cat# 80376) were filled with pre-warmed TeSR-E6. PT-EKO were excised using a microsurgical blade from Transwell membranes, with a small square of membrane remaining, and placed tissue side-up onto each chamber. Chips were then sealed with the adhesive Ibidi Polymer Coverslip whilst avoiding bubble formation. Each chip was connected to a syringe pump (New Era, Farmingdale, NY, USA, cat# 300) using Tygon tubing (Biorad, CA, USA, cat# 7318215) of equal lengths and a 1ml syringe (Terumo, Leuven, Belgium, cat# SS+01T) containing TeSR-E6. After 48 hours, PT-EKO cultured on static Transwell and OoC platforms were harvested and fixed with 4% paraformaldehyde for immunofluorescence staining.

### Quantification of mitochondrial distribution

To analyse the distribution of TOMM20, Fiji Image J was used to generate binary masks for the LTL, EPCAM and TOMM20 fluorescence channels by applying intensity-based thresholding. LTL and EPCAM masks were then used to define regions of interest for TOMM20 quantification, with the Boolean operation “and” applied between masks to distinguish colocalised TOMM20/LTL signal versus colocalised TOMM20/EPCAM signal. The resulting masks were then analysed to express results in pixels.

### Cryopreservation and thawing of D13 PT-EKO monolayer progenitors

PT-EKO progenitors at D13 of monolayer differentiation were dissociated using TrypLE Select Enzyme (1X; Gibco, ThermoFisher Scientific, MA, USA, cat# 1256029) following the same procedure as for organoid generation above. After stopping the enzymatic reaction in 2% FBS (Gibco, ThermoFisher Scientific, MA, USA) in TeSR-E6 (STEMCELL Technologies, Vancouver, Canda), cell counts were performed using Countess™ 3 FL Automated Cell Counter (Invitrogen, ThermoFisher Scientific, MA, USA) and cells were centrifuged (300g for 3 mins at room temperature). Cell pellets were resuspended in ice-cold Cryo-SFM freezing medium (Promo-Cell, Heidelberg, Germany, cat# C-29912) at a density of approximately 6 x10^6^ cells/mL before transferring 1mL aliquots into cryovials (Corning, New York, USA). Cryovials were frozen overnight at -80°C in Mr Frosty (Nalgene, New York, USA) before transferring to liquid nitrogen.

For the generation of PT-EKO from cryopreserved D13 monolayers, cryovials were warmed in a 37°C waterbath until a small piece of ice still remained. In the biosafety cabinet, thawed cells were transferred in a dropwise-manner to a 15mL tube containing cold 2% FBS (Gibco) in TeSR-E6 (STEMCELL Technologies) supplemented with 10µM ROCKi (TOCRIS Bioscience, Bristol, UK) and centrifuged at 300g (3mins, room temperature) to remove freezing medium. The cell pellets were resuspended in a defined volume of 2%FBS (Gibco) in TeSR-E6 (STEMCELL Technologies) supplemented with 10µM ROCKi (TOCRIS Bioscience), cell counts performed, and the total volume adjusted to 350,000 cells per 200uL (1 organoid equivalent per 200µL aliquot). After aliquoting 200uL per 1.5mL tube (Axygen, Corning, NY, USA) for each organoid to be generated, micromasses were generated via 3X centrifugations and transferred to Transwell membranes as above as previously published ^43^ with minimal changes except for the supplementation of CHIR-pulse media (TeSR-E6 containing 5µM CHIR [TOCRIS Bioscience]) with10µM ROCKi (TOCRIS Bioscience). Following the CHIR pluse, media was replaced with TeSR-E6 containing 200ng/mL FGF9 (R&D Systems, Minneapolis, MN, USA) as per published protocols ^43^.

### Fatty acid-modified culture conditions

D13+8 PT-EKOs were incubated for 2 days in a 1:1 mixture of standard culture media (TeSR-E6) and fatty acid-modified media as detailed below, before replacing with 100% fatty acid-modified media for a further 4 days of culture and refreshing every second day prior to harvesting. Short chain fatty acid-modified media consisted of glucose-free and glutamine-free DMEM (Gibco, ThermoFisher, cat# A1443001) supplemented with 0.5 mM GlutaMAX (Gibco, ThermoFisher, cat# 35050061), 1% penicillin/streptomycin, 5mM glucose, 2 mM sodium acetate (Sigma Aldrich, cat# S5636), and 500 µM sodium butyrate (Sigma Aldrich, cat# 303410). Long chain fatty acid-modified media consisted of glucose-free and glutamine-free DMEM (Gibco, ThermoFisher, cat# A1443001) supplemented with 0.5 mM GlutaMAX (Gibco, ThermoFisher, cat# 35050061), 1% penicillin/streptomycin, 5 mM glucose, 2% FCS, and 200 µM sodium palmitate (Sigma cat# P9767; conjugated with 33 µM fatty-acid-free BSA [Sigma cat# A7030] at a 6:1 ratio).

### Fatty acid oxidation assay

FAO-Glo^TM^ Fatty Acid Oxidation Assay (Promega, USA, cat# CS3271A08) was used to measure fatty acid oxidation via CPT1a in organoids, according to manufacturer’s protocol. Briefly, organoids of age D13+14 - D13+16 were excised from their Transwell membranes and transferred to a 96-well plate (Corning cat# CLS3917) before 100uL of FAO substrate was added per well. After 2 hours of incubation at 37⁰C, 100uL of FAO Detection Reagent was added followed by a further 10 minute incubation. Luciferase readings were measured immediately thereafter on a Tecan Spark® plate reader operated using the Tecan Spark Control software v3.2 SP1. Luminescence signals were normalised to control wells without cells. For inhibitor-treated conditions, organoids were pre-incubated in 20uM of Etomoxir (ETO; MedChemExpress, USA, cat# HY-50202) for 2, 4, or 24 hours prior to removal from the Transwells for FAO substrate addition and luciferase readings.

### FACS isolation and expansion of PT cells from PT-EKO

D13+14 PT-EKO were transferred to a 6cm dish (Greiner Bio-One, Kremsmünster, Austria) and manually dissociated for 1 minute using scalpel followed by addition of 1mL (per 24 organoids) of 1:1 TrypLE Select (Gibco) and Accutase (BD Biosciences, New Jersey, USA) enzyme mixture. The dish was incubated at 37°C 15 mins, mechanically pipetting every 3 minutes using a P1000 pipette. After 15 minutes, the enzymatic reaction was stopped by adding 1mL of 2% FBS (Gibco) in TeSR-E6 (STEMCELL Technologies) and the cell suspension passed through a 40µM cell strainer (Falcon, Corning, New York, USA) followed by a round bottom FACS tube with cell strainer cap (BD Biosciences, NJ, USA). Cells were centrifuged at 300g for 3mins at room temperature and the cell pellet resuspended in 500µL of cold 2%FBS in Hanks’ Balanced Salt Solution (HBSS, Gibco).

After cell counting, the density was adjusted to a range of 1.0 x 10^6^ - 5 x 10^6^ cells/mL for primary antibody labeling with 20µg/mL biotinylated *Lotus tetragonobulus* lectin (LTL, Vector Laboratories, California, USA). Cells were incubated in primary antibody on ice for 15 minutes then washed by diluting with 3X volume of 2% FBS/HBSS (FACS wash) followed by centrifugation (300g for 3 minutes at room temperature). The supernatant was aspirated and the cell pellet was resuspended in 1:300 streptavidin-conjugated Alexa Fluor-647 secondary antibody (Invitrogen, ThermoFisher Scientific, MA, USA) and 1:300 anti-EpCAM-conjugated Alexa Fluor-488 (Biolegend, California, USA) in FACS wash. Following 15 minutes incubation on ice and washing as per above, LTL+ EpCAM+ cells were isolated using the FACSAria Fusion (BD Biosciences). Sorted cells were collected in 2% FBS/TeSR-E6.

Isolated PT cells were maintained and expanded according to methods published previously with adaptions ^12^. Briefly, 96 well plastic plates (NUNC, ThermoFisher Scientific, MA, USA) or 96 well glass-bottom plates (Cellvis, California, USA) were coated with 5µg/cm^2^ collagen IV (Sigma) or 200µL of 20µg/mL Geltrex (Gibco), respectively, and incubated for 1 hour (collagen IV) or 3 hours (Geltrex) at 37°C. Prior to seeding, wells were washed 3 times with PBS. Isolated PT cells were resuspended in PT media (below) supplemented overnight with 10µM ROCK inhibitor (ROCKi, Y-27632; TOCRIS Bioscience) to promote cell survival following FACS and seeded into coated plates at a density of 300,000 cells/cm^2^ (PT media: DMEM low glucose [Gibco], Ham’s F12 nutrient mix [Gibco], 5mM HEPES [Gibco], penicillin-streptomycin [Gibco], 50µM ascorbic acid [Sigma, Millipore Sigma, Burlington, Massachusetts, USA], ITS [0.86µM insulin, 0.06µM transferrin, 0.06µM selenium; Sigma], 10ng/mL epidermal growth factor [EGF; Millipore, Sigma, Burlington, Massachusetts, USA], 25ng/mL hydrocortisone [Sigma], 10ng/mL 3,3’-Triiodo-L-thyronine [T3; Sigma], 10µM TGF-β inhibitor SB431542 [STEMCELL Technologies], and 1µM TGF-β inhibitor A83-01 [TOCRIS Bioscience, Bristol, UK]). The following day, the PT media + 10µM ROCKi was replaced with PT media alone and media changed every 2 days.

Upon reaching 80% - 90% confluency, cells were passaged by washing with PBS followed by 0.5mM EDTA (Gibco). The cells were then incubated in 200µL of TrypLE (Gibco) at 37°C for 7-9 minutes before gently pipetting up and down 3 – 5 times to detach. The detached PT cells were collected in a 1.5 mL tube containing 0.5mL of 10% FBS / PBS solution and centrifuged at 300g for 5 minutes (room temperature). After gently aspirating the supernatant, the cells were resuspended in PT media supplemented with 10µM ROCKi (TOCRIS Bioscience) and divided at a passaging ratio of 1:2 between wells of fresh collagen IV-coated (plastic) or Geltrex-coated (glass) plates.

### Functional uptake assays in isolated and primary PT cells

Isolated PT cells (cultured as per above) and hRPTECs (Lonza Bioscience, Maryland, USA), cultured as per manufacturer instructions) were expanded to passage 2-3 and seeded into glass-bottomed 96-well plates coated with 20µg/mL Geltrex at a density of 100,000 cells/cm^2^, allowing for 3 replicate wells per test and control condition for each substrate (TRITC-Albumin substrate [10 mg/mL stock, Sigma Aldrich], 4-(4-(dimethylamino)styryl)-N-methylpyridinium substrate [ASP; 1mM stock, ], and 4’,6-diamindino-2-phenylindole substrate [DAPI; 5mg/mL stock, Thermo Fisher Scientific]). Once confluent (approximately 48 hours after seeding), culture media was firstly aspirated from the control wells, replaced with 200µL per well of HBSS, and incubated for 10 minutes (37 °C, 5% CO2 and 5% O2; used for all incubation steps throughout uptake assays). After 10 minutes, culture media / HBSS was aspirated from all wells and replaced with either plain HBSS (TRITC-albumin uptake control and test wells to commence washing) or HBSS containing 1mM Cimetidine (Sigma Aldrich) (ASP and DAPI control wells to commence transporter inhibition). After 30 minutes incubation, Cimetidine solution was aspirated from the ASP and DAPI control wells and replaced with 200µL per well of substrate + inhibitor solution, with the addition of DRAQ7 (Thermo Fisher Scientific) to the DAPI condition to label dead cells (ASP control = 2.5µM ASP + 1mM Cimetidine in PBS. DAPI control = 1:6000 DAPI + 1mM Cimetidine + 1:1000 DRAQ7 in HBSS). In parallel, substrate-only solutions were added to the test wells (ASP test = 2.5µM ASP in HBSS. DAPI test = 1:6000 DAPI + 1:1000 DRAQ7 in HBSS. Albumin test = 1:1000 TRITC-Albumin in HBSS). TRITC-albumin control wells remained in HBSS. The plate was incubated for 15 minutes before 3 quick washes in HBSS and imaging immediately using a ZEISS LSM 780 confocal microscope. Live confocal images were segmented using Fiji ImageJ 1.54p Labkit plugin (Version 0.4.0). For each condition, classifiers were trained for the detection of TRITC (568nm channel), ASP (488nm channel), and DAPI (405nm channel) with DRAQ7 (647nm channel). However, negligible overlap was present between DAPI and DRAQ7. A custom macro was developed to batch analyse samples across conditions. The area occupied by fluorescent pixels was quantified and expressed as a percentage of the total image area (taken at equivalent magnification, zoom, and intensity settings for test and controls within each condition).

## AUTHOR CONTRIBUTIONS

Conceptualization: J.M.V., M.H.L.; Funding Acquisition: M.H.L., J.M.V.; Project Administration: J.M.V.; Data Curation: S.M., K.S.T., J.M.V., S.B.W., K.T.L., E.I.S.; Investigation: S.M., K.S.T., M.C, S.B.W., J.M.V.; Formal Analysis: S.M., K.S.T., S.B.W., K.T.L., J.M.V., B.G., E.I.S.; Methodology: J.M.V., S.M., K.S.T., S.B.W., M.C., R.J.M.; Resources: R.J.M, M.C.; Validation: S.M., K.S.T., J.M.V, S.B.W.; Visualization: J.M.V., S.M., K.S.T., K.T.L, S.B.W., B.G.; Software: S.B.W, K.T.L, E.I.S., J.M.V.; Supervision: J.M.V., M.H.L., R.J.M., K.T.L.; Writing: Original Draft: J.M.V.; Writing – Review and Editing: J.M.V., M.H.L., S.M., K.S.T., S.B.W., R.J.M.

## ACKNOWLEDGEMENTS

We acknowledge MCRI Operational Infrastructure Support and the Stafford Fox Medical Research Foundation MCRI Genome Editing Facility for the generation of pluripotent stem cell lines; Matthew Burton and the Murdoch Children’s Research Institute Microscopy Core. We also thank Professor William Fissell and Rachel C Evans (Vanderbilt University, NA, USA) for their advice on proximal tubule cell culture. Some schematics contain images adapted from BioRender.com. We acknowledge the research reported in this publication was supported by The Novo Nordisk Foundation Center for Stem Cell Medicine (*reNEW*) [NNF21CC0073729], the Medical Research Future Fund (MRFF2040599), and The University of Melbourne, Melbourne Medical School. For the purposes of open access, the author has applied a CC BY public copyright license to any Author Accepted Manuscript version arising from this submission.

## DATA AND CODE AVAILABILITY

ScRNAseq analyses in the current study were performed on our existing published single cell dataset of PT-enhanced organoids (GEO accession: GSE184928). Bulk RNA sequencing datasets have been uploaded to GEO and will be made publicly available upon publication (GEO accession: GSE346993). Code for the scRNAseq and bulk RNAseq analyses in the current study is available on request and will be made publicly available upon publication in the Kidney Regeneration Github repository (https://github.com/KidneyRegeneration/Vanslambrouck2026).

