## Supplementary Data for "Proximal tubule-enhanced kidney organoids with S3-like cell enrichment model physiological responses"

***Equal first author contribution**

**#Equal last author contribution**

**Running title:** S3-enriched proximal tubule model from stem cells

#### SUPPLEMENTARY TABLES

**Supplementary Table 1:** Human nomenclature of PT segment markers curated from literature detailing mammalian distribution, prioritizing human data where available.

| PT marker (gene) | Corresponding protein | Reported PT distribution | References |
|---|---|---|---|
| SLC6A19 | Sodium-dependent neutral amino acid transporter B(0)AT1 / SLC6A19 | S1 | 121 |
| SLC36A2 | Protein-coupled amino acid transporter 2 (PAT2 / SLC26A2) | S1 | 121 |
| RBP4 | Retinol-binding protein 4 (RBP4) | S1>S2 | 122 |
| SLC2A2 | Glucose transporter 2 (GLUT2) | S1>S2 | 46,47 |
| SLC5A2 | Sodium-glucose co-transporter 2 (SGLT2) | S1>S2 | 46-49 |
| SLC34A1 | Sodium-dependent phosphate transport protein 2A (NaPi-2a) | S1>S2 | 123 |
| ANPEP | Aminopeptidase N (CD13) | S1>S3 | 49,124 |
| GPX3 | Glutathione peroxidate (GPX3) | S1>S3 | 125 |
| CUBN | Cubilin (CUBILIN / CUBN) | S1>S3 | 122 |
| SLC47A1 | Multidrug and toxin extrusion protein 1 (MATE1) | S1>S3 | 93 |
| SLC7A9 | Cystine / glutamate transporter (b(0,+)AT1 / SLC7A9) | S1>S3 | 92 |
| SLC22A8 | Organic anion transporter 3 (OAT3) | S1>S3 | 126 |
| AQP1 | Aquaporin 1 (AQP1) | S1-S3 | 125 |
| CA4 | Carbonic anhydrase 4 (CA4) | S2 | 127 |
| LYZ | Lysozyme C (LYZ) | S2 | 122 |
| SLC6A20 | System IMINO transporter (XT3 / SIT1 / SLC6A20) | S2-S3 | 121 |
| SLC15A2 | Peptide transporter 2 (PEPT2) | S2-S3 | 128 |
| SLC22A10 | Organic anion transporter 5 (OAT5) | S2-S3 | 129 |
| SLC5A1 | Sodium-glucose co-transporter 1 (SGLT1) | S2<S3 | 46-49 |
| ALPI | Intestinal-type alkaline phosphatase (IAP) | S2<S3 | 130,131 |
| AGT | Angiotensinogen (AGT) | S2<S3 | 92 |
| SLC2A1 | Glucose transporter 1 (GLUT1) | S2<S3 | 46,47 |
| SLC1A1 | Excitatory amino acid transporter 3 (EAAT3 / SLC1A1) | S2<S3 | 92 |
| SLC39A8 | Zinc transporter ZIP8 | S1<S3 | 132 |
| SLC39A14 | Zinc transporter ZIP14 | S1<S3 | 132 |
| SLC38A3 | Sodium-coupled neutral amino acid transporter 3 (SNAT3) | S1<S3 | 132 |

**Supplementary Table 2:** Average PT isolation details based on 4 independent experiments of PT-EKOs (starting cell number: 250,000 cells per micromass) that displayed typical morphology, harvested D13+14 – D14+19.

| Average total cell number obtained per organoid post-dissociation | Average PT cell number isolated per organoid | Average PT cell yield per 500,000 cells (dissociated PT-EKO) | Average FACS PT proportion (% of live single cells [parent]) |
|---|---|---|---|
| 440,309 (min. 318, 056, max. 594,907) | 25,174 (min. 12,372, max. 34,074) | 28,093 (min. 18,772, max. 32,751) | 17.3% (min. 12.0%, max. 20.3%) |

### SUPPLEMENTARY FIGURES

#### Supplementary Figure 1

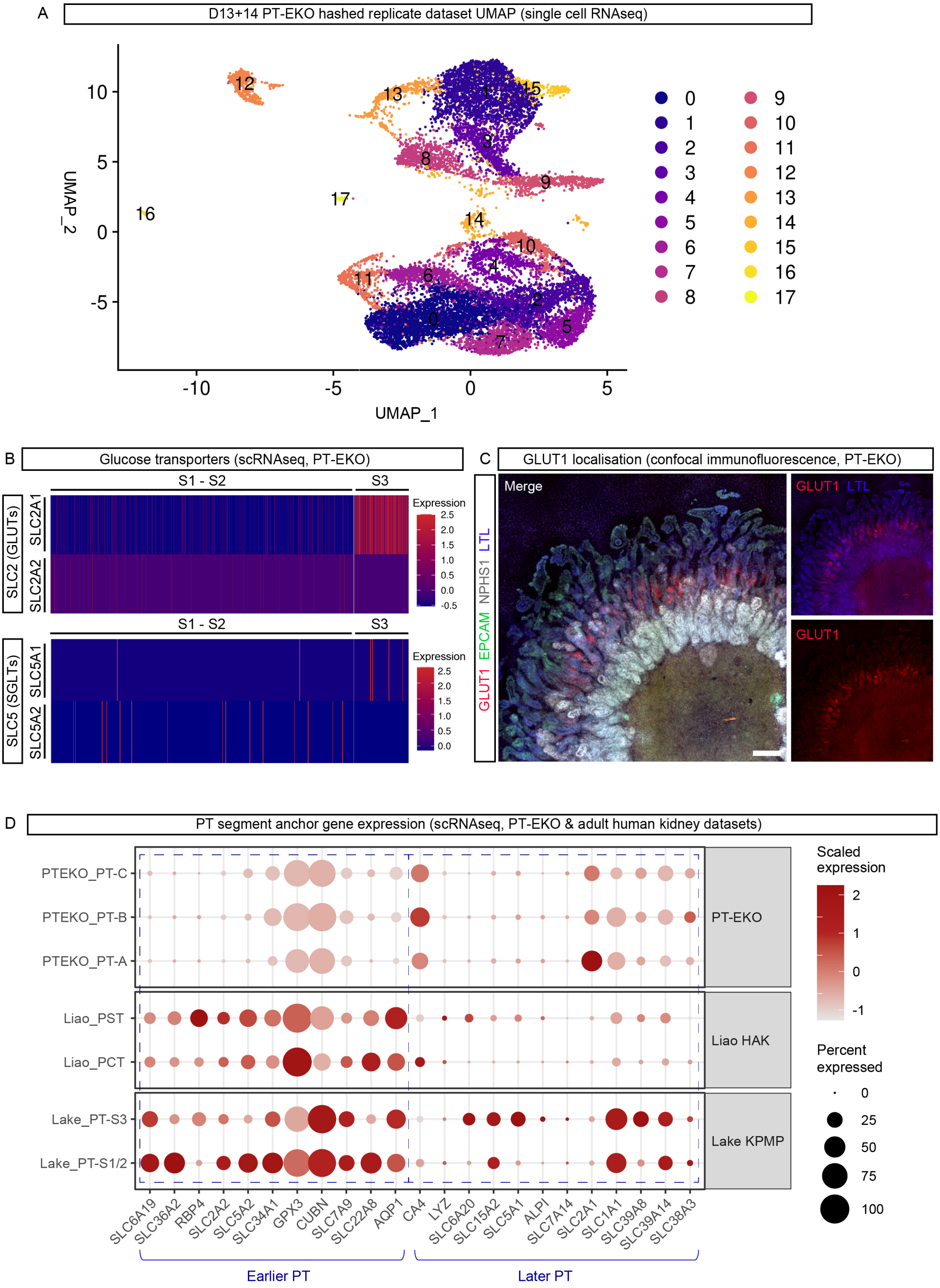

30 **Supplementary Figure 2**

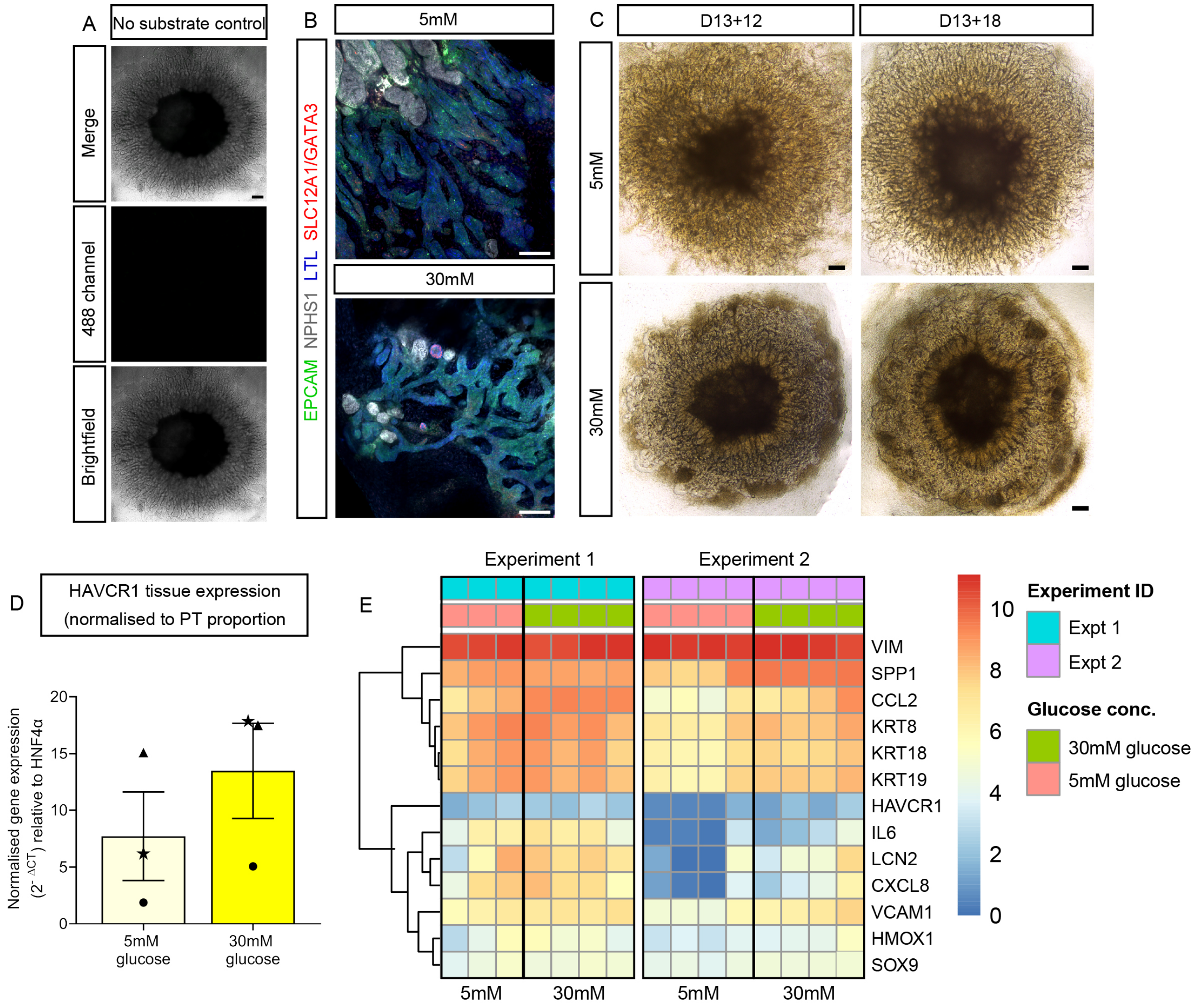


31

#### Supplementary Figure 3

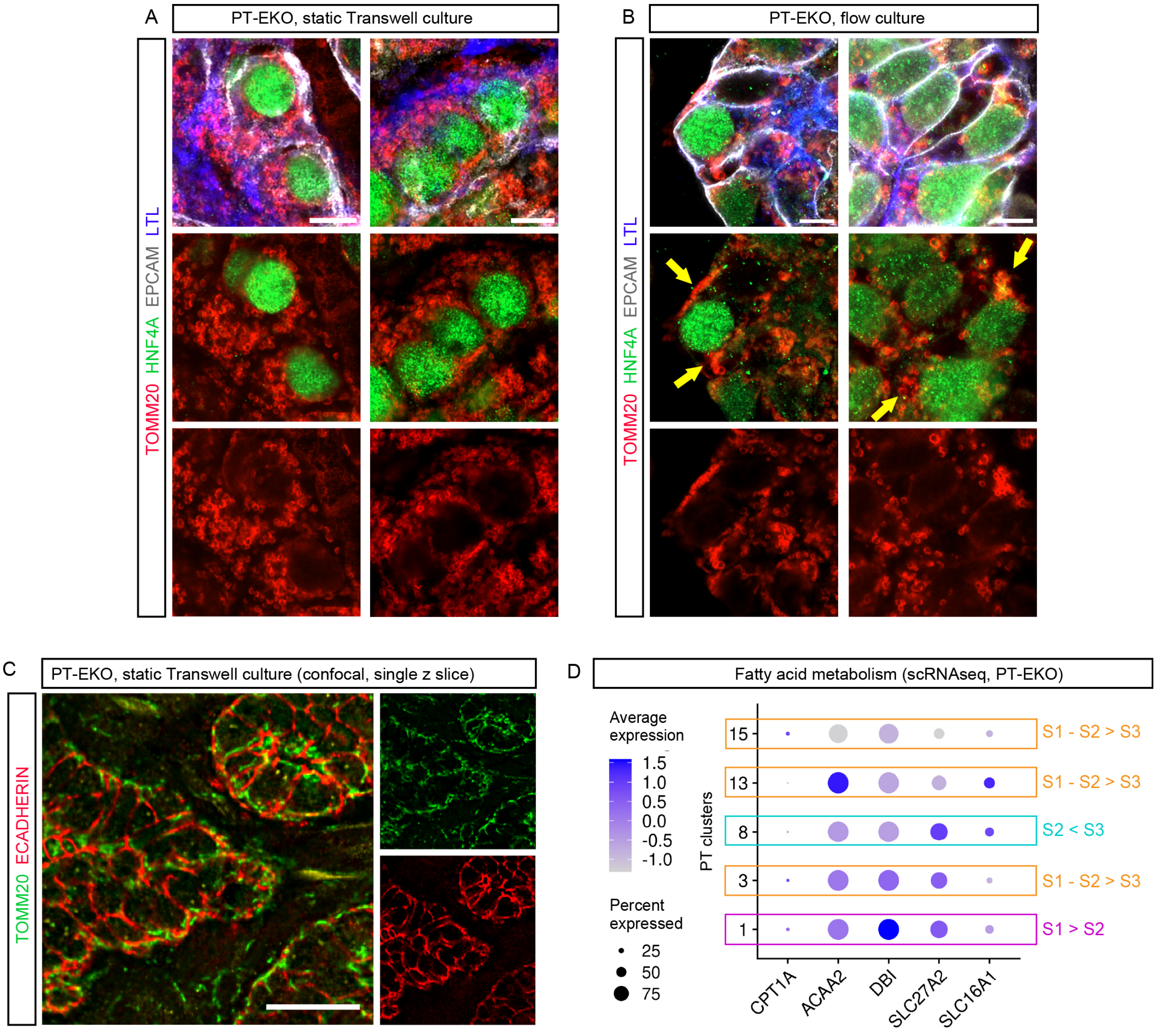

### Supplementary Figure 4

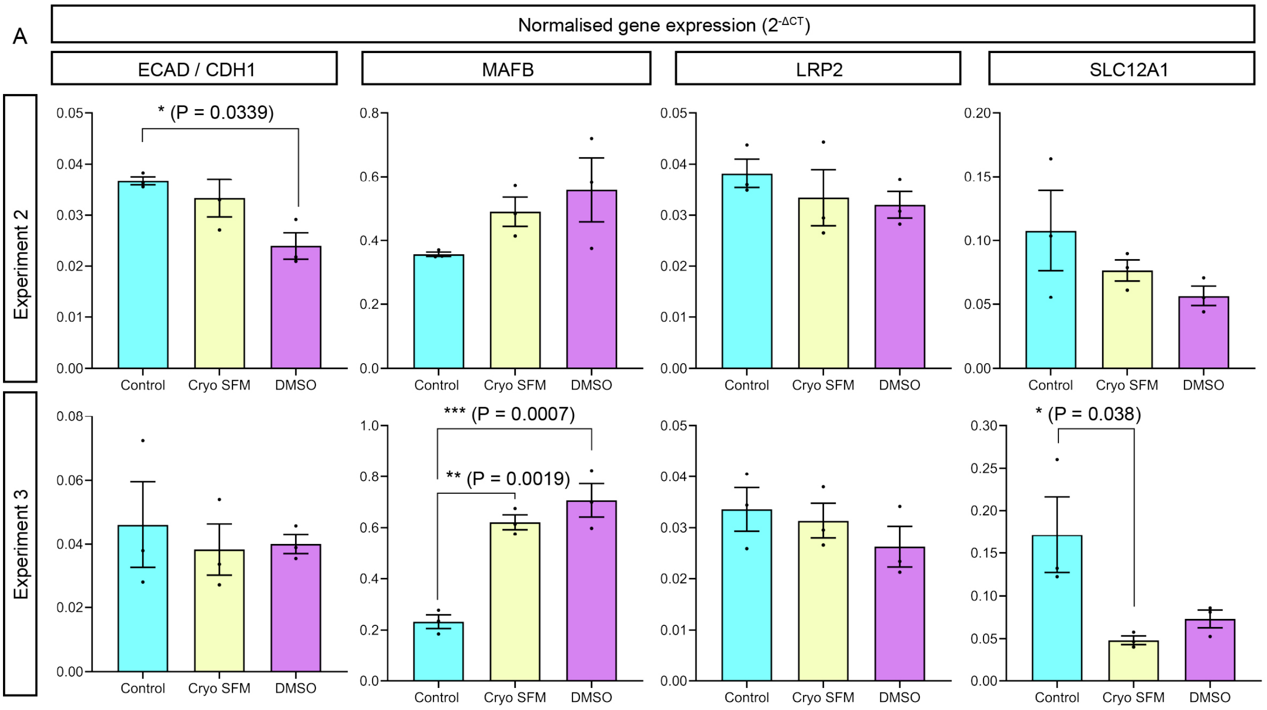

#### Supplementary Figure 5

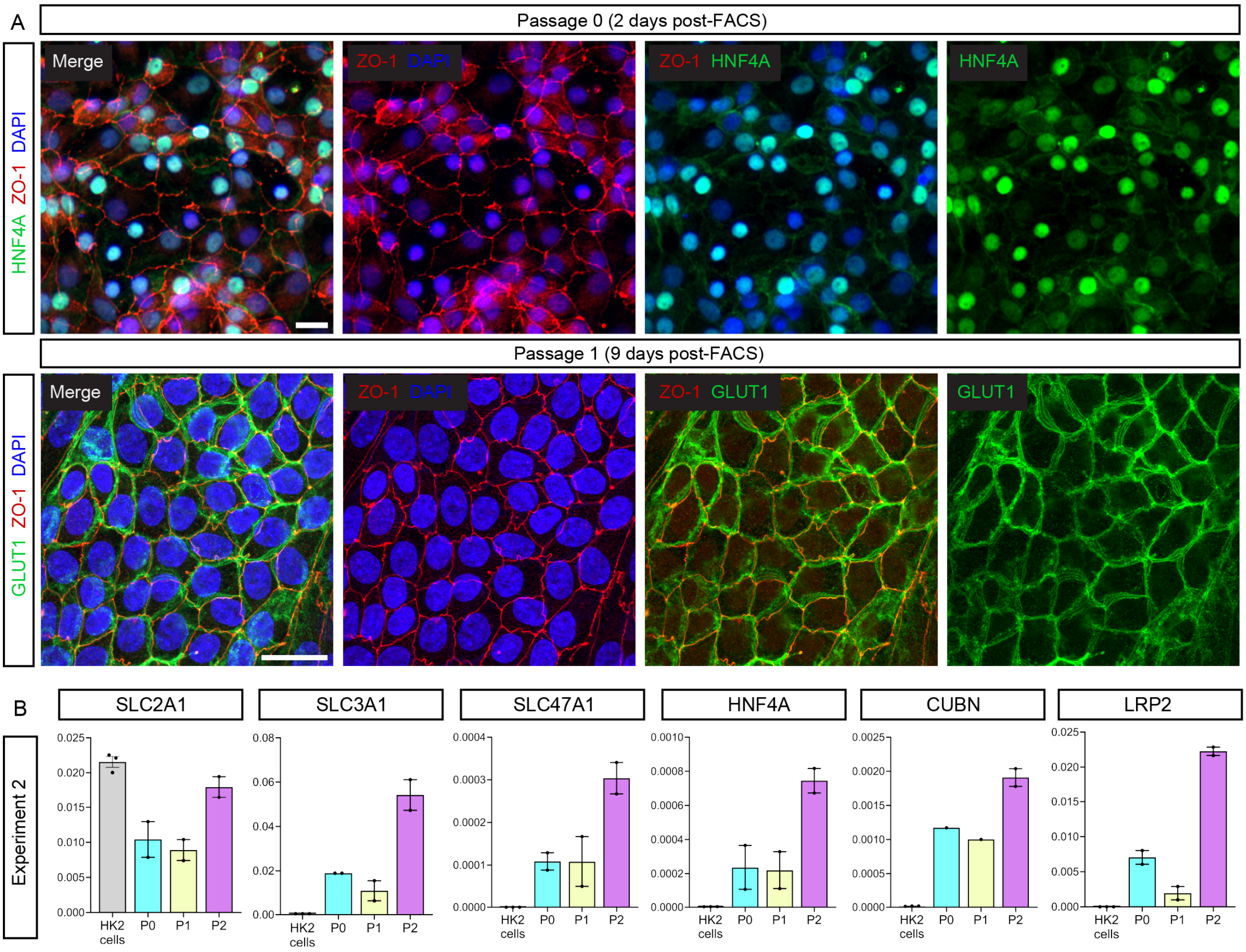

### Supplementary Figure 6

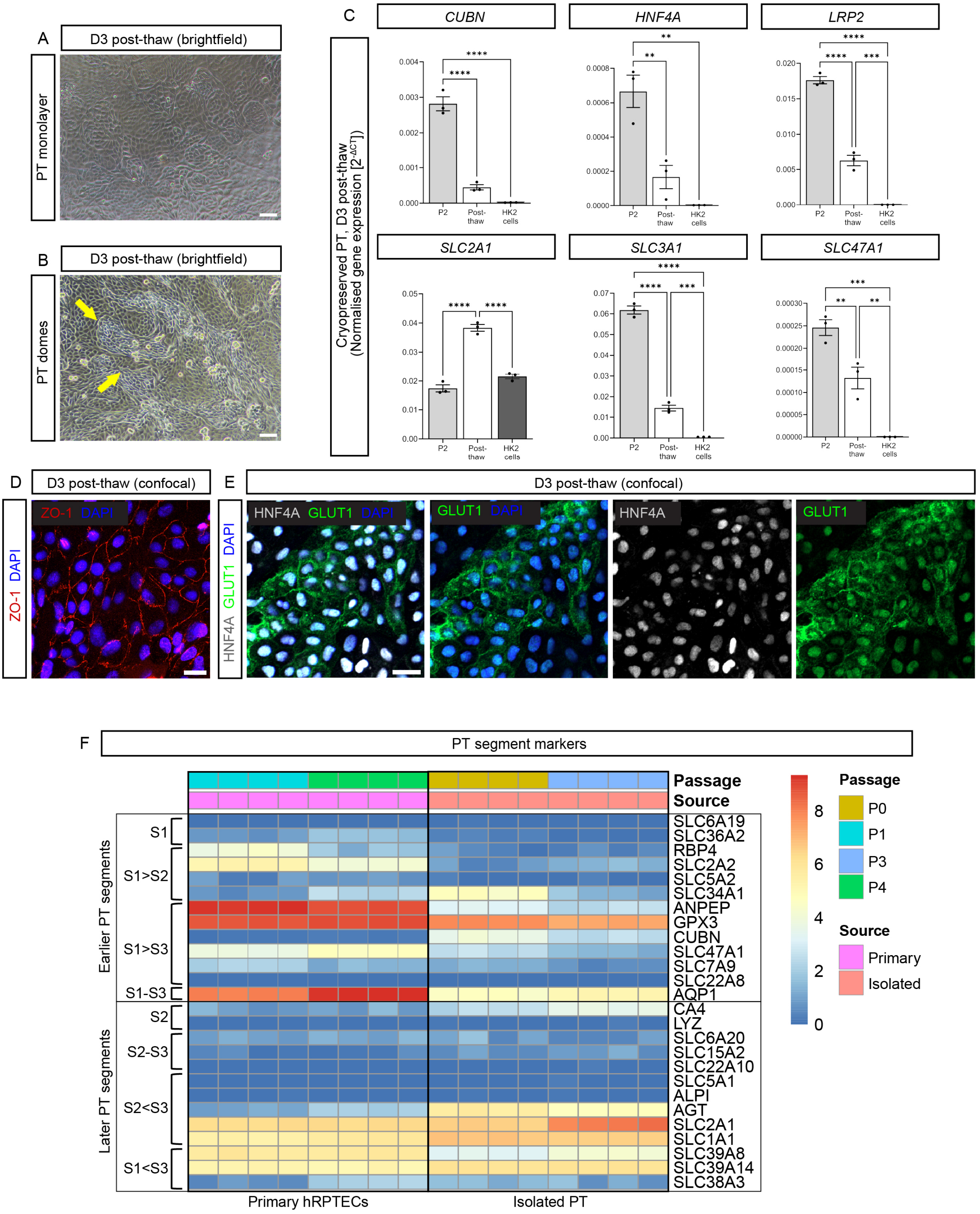

#### SUPPLEMENTARY FIGURE LEGENDS

**Supplementary Figure 1: PT-EKO show evidence of enrichment for S3-like cell types, PT physiological characteristics, and can be generated from cryopreserved D13 monolayers. A**. UMAP plot of D13+14 PT-EKO samples (pooled replicates) from Vanslambrouck *et al.* (2022) [41] highlighting 17 distinct cell clusters and showing clusters 1, 3, 8, 13, and 15 identified as PT populations in the original analysis. **B.** Heatmap showing the expression of glucose transporters with known differential expression patterns in PT segments (S1-S2 or S3). S1-S2 represents merged clusters 1, 3, 13, 15, while S3 represents cluster 8. **C**. Confocal immunofluorescence of PT-EKO stained with markers of PT brush border membrane, S1-S3 (*Lotus tetragonolobus* lectin [LTL]; blue), proximal tubule S2<S3 basolateral glucose transporter (GLUT1; red), podocytes of the glomeruli (NEPHRIN [NPHS1]; grey), and distal nephron epithelium (E-CADHERIN [ECAD]; green). Scale bar represents 200µm. **D.** Dot plot depicting the expression of PT segment anchor genes curated from literature (Supplementary Table 1) used further validate PT cluster identity and merge similar populations in Figure 1D. Datasets represent PT-EKO S3-like (PTEKO_PT-A [clusters 8 and 3]), PT-EKO S1-S2-like (PTEKO_PT-B [cluster 15] and PTEKO_PT-C [clusters 1 and 13]), adult human proximal convoluted / S1-S2 (Liao_PCT and Lake_PT-S1/2), and adult human proximal straight (Liao_PST and Lake_PT-S3) [52,53]. Dot colour and size in dot plots represent scaled gene expression and percentage of cells expressing each gene, respectively.

**Supplementary Figure 2: Glucose concentration influences morphology and injury marker expression in PT-EKO. A**. No substrate control PT-EKO (without 2-NBDG exposure) confirming lack of green fluorescence in the 488 channel (middle image). Top image shows brightfield and 488 channel overlay. Scale bar represents 200 µm. **B**. Confocal immunofluorescence images of D13+18 PT-EKOs cultured in 5mM or 30mM glucose stained for markers of PT (LTL; blue), podocytes of the glomeruli (NPHS1; grey), nephron epithelium (EPCAM; green), and distal tubule (loop of Henle thick ascending limb [TAL, SLC12A1, apical] and connecting segment [GATA3, nuclear]; red). Scale bars represent 100µm.**C.** Brightfield images of PT-EKO at D13+12 and D13+18 following exposure to glucose-modified media containing 5mM or 30mM glucose. Scale bars represent 200µm. **D.** qRT-PCR of Kidney Injury Molecule–1 (KIM-1) gene, *HAVCRI*, in D13+18 PT-EKOs exposure to 5mM (light yellow bar) and 30mM (dark yellow bar) glucose, with expression values normalised to the expression of housekeeping gene *GAPDH* and compensating for differences in PT proportion (expressed as a ratio of *HNF4A*). Error bars indicate SEM from n = 3 independent experiments, identified by datapoint symbol, each datapoint representing the average of n= 4 biological replicates. **E.** Heatmap showing the expression of a curated list of kidney injury markers in PT-EKO exposed to 5mM or 30mM glucose-containing media (n = 3 – 4 biological replicates per condition across 2 independent replicate experiments). Gene expression depicted as log-transformed transcripts per million.

**Supplementary Figure 3: PT-EKO display characteristic metabolic features. A-B.** High magnification examples of confocal immunofluorescence images from Figure 3AB, depicting D13+18 PT-EKO cultured in standard static conditions on Transwell membranes (A) and under 20µL/hour media flow within Ibidi 3D Perfusion Slides (B), showing mitochondria (TOMM20; red), PT (LTL; blue, HNF4A; green), and nephron epithelium (EPCAM; grey). Scale bars represent 10µm. **C.** Confocal immunofluorescence of PT-EKO cultured in static Transwell conditions showing TOMM20+ mitochondria (green) within E-CADHERIN+ distal tubules (red). Scale bar represents 20µm. **D.** Dot plot depicting the expression of fatty acid metabolism genes in PT clusters within the D13+14 PT-EKO dataset (4 hashed replicates merged). Dot colour and size in dot plots represent unscaled gene expression and percentage of cells expressing each gene, respectively.

**Supplementary Figure 4: Cryopreserved D13 monolayers produce PT-EKO with retained proximal nephron gene expression. A.** qRT-PCR from experiment in Figure 4D depicting an additional 2 independent replicate experiments, comparing nephron markers (MAFB; podocytes, LRP2; PT, SLC12A1; loop of Henle TAL, and ECAD; distal nephron) in D13+14 control PT-EKO (cyan bars), as well as PT-EKO generated from Cryo-SFM (yellow bars) and DMSO (magenta bars) cryopreserved D13 monolayers. Error bars indicate SEM from n = 3 biological replicates per experiment. Statistical significance was assessed using a one-way ANOVA with Tukey's multiple comparisons test. Asterisks denote two-tailed P values of $\leq 0.05$ (*), $\leq 0.01$ (**), and $\leq 0.001$ (***) for pairwise comparison.

**Supplementary Figure 5: Isolated PT cells from PT-EKO show expected protein and gene expression across multiple passages. A.** Confocal immunofluorescence images of isolated PT cultures at 2- (passage 0) and 9-days (passage 1) post-FACS, depicting tight junction protein expression (ZO-1; red), nuclear transcription factor HNF4A (green, top row), membrane-bound glucose transporter expression (GLUT1; green, bottom row), and cell nuclei (DAPI; blue). Scale bars represent 20µm. **B.** qRT-PCR of PT genes in isolated PT cells across a 7 day time course (passage 0 [P0; cyan bars], passage 1 [P1; yellow bars], passage 2 [P2; magenta bars]) compared to HK2 cells (grey bars). Error bars represent SEM of n = 2 biological replicates.

**Supplementary Figure 6: Isolated PT cells are amenable to cryopreservation and show an S3-like identity. A-B.** Brightfield images of cryopreserved PT cells at 3 days post-thaw, depicting cuboidal morphology (A) and the formation of domes in monolayer culture (yellow arrows; B). Scale bars represent 100µm. **C.** qRT-PCR of PT marker genes in PT-EKO-derived cryopreserved PT cells 3 days post-thaw (white bars) alongside PT cells prior to cryopreservation (passage 2 [P2]; light grey bars) and commercial HK2 cell cultures (dark grey bars). Error bars represent SEM of n = 3 biological replicates. Statistical significance was assessed using a one-way ANOVA with Tukey's multiple comparisons test. Asterisks denote two-tailed P values of ≤0.05 (*), ≤0.01 (**), ≤0.0001 (***), and ≤ 0.0001 (****) for pairwise comparisons. **D-E.** Confocal immunofluorescence images of isolated cryopreserved PT cultures at 3 days post-thaw depicting tight junction protein expression (ZO-1; red), nuclear transcription factor HNF4A (grey), membrane-bound glucose transporter expression (GLUT1; green), and cell nuclei (DAPI; blue). Scale bars represent 20µm. **F**. Heatmap showing gene expression levels for markers of early (S1-S2) and late (S2-S3) PT segments (also refer to Supplementary Table 1) within primary hRPTECs and isolated PT cells across 2 passages with n = 4 replicate wells per condition. Gene expression depicted as log-transformed transcripts per million.
